# 3D Imaging Reveals Changes in the Neurovascular Architecture of the Murine Calvarium with Aging

**DOI:** 10.1101/2024.03.28.587299

**Authors:** Allison L. Horenberg, Yunke Ren, Alexandra N. Rindone, Arvind P. Pathak, Warren L. Grayson

## Abstract

Calvarial nerves, along with vasculature, influence skull formation during development and following injury, but it remains unclear how calvarial nerves are spatially distributed during postnatal growth and aging. Studying the spatial distribution of nerves in the skull remains challenging due to a lack of methods to image and quantify 3D structures in intact bone. To visualize calvarial 3D neurovascular architecture, we imaged nerves and endothelial cells with lightsheet microscopy. We employed machine-learning-based segmentation to facilitate high-resolution characterization from post-natal day 0 (P0) to Week 80 (80wk). We found that TUBB3+ nerve density decreased with aging with the frontal bone demonstrating earlier onset age-related nerve loss than the parietal bone. In addition, nerves in the periosteum and dura mater exhibited similar yet distinct temporal patterns of nerve growth and loss. While no difference was observed in TUBB3+ nerves during skeletal maturation (P0 ⟶ 12wk), we did observe an increase in the volume of unmyelinated nerves in the dura mater. Regarding calvarial vasculature, larger CD31^hi^Emcn^-^ vessel density increased with aging, while CD31^hi^Emcn^hi^ vessel density was reduced. For all nerve markers studied, calvarial nerves maintained a preferential spatial association with CD31^hi^Emcn^hi^ vessels that decreased with aging. Additionally, we used a model of Apert syndrome that demonstrates early coronal suture fusion to explore the impact of suture-related disease on neurovascular architecture. We identified a mild dysregulation of dural nerves and minor shifts in vessel populations. Collectively, this 3D, spatiotemporal characterization of calvarial nerves throughout the lifespan and provides new insights into age-induced neurovascular architecture.

## Introduction

A dense network of peripheral nerves reside throughout bone and have known pro-osteogenic roles during development and after injury^1^. Following initial ossification *in utero*, murine bones continue to develop, then remodel throughout adulthood. Ultimately, their capacity to effectively remodel and regenerate diminishes, and this age-dependent loss is thought to be primarily mediated by cellular senescence.^2^ Osteoprogenitor cells reduce in number and proliferative capacity with aging, resulting in a loss of bone formation and increased bone fragility.^3,4^ Other cell types in the bone microenvironment also mediate bone loss with aging. In aged fracture healing studies, macrophages maintain a pro-inflammatory phenotype and exhibit reduced bone induction capacity compared to macrophages in young mice.^5^ Likewise, in aged long bones, Type H (CD31^hi^Emcn^hi^) blood vessel density declines, resulting in decreased perfusion, increased hypoxia and reactive oxygen species production, and significantly lower levels of hematopoietic factors.^6^ While these cell types have known roles in modulating bone structure throughout the lifespan, the peripheral nerves have not been implicated in mediating changes in bone structure during aging.

Bone contains sensory nerves, which generate the pain associated with headaches and bone fracture, as well as sympathetic nerves.^7–10^ Apart from skeletal pain, sensory nerves play roles in successful healing of bone fracture injuries. Additionally, both nerve subtypes are closely linked with blood vessels and play a role in their growth and differentiation.^11–13^ In bone, nerves are found in close vicinity of blood vessels, both in homeostasis and in bone fracture.^8–10,14–21^ CD31^hi^Endomucin (Emcn)^hi^ blood vessels are a vascular subtype that is crucial for bone homeostasis, development, and response to injury, however, they have yet to be implicated in nerve signaling pathways.^22,23^ In addition to aging models, the neurovascular architecture in an FGFR-related craniosynostosis syndrome, characterized by premature suture closure is of interest, as previous studies have found that inhibiting nerves through TrkA inhibition can lead to enhanced sutural closure and dysregulation of skeletal stem cell proliferation.^24^ Craniosynostosis syndromes are characterized by suture fusion early in development. A majority of craniosynostosis syndromes are associated with mutations in the FGFR2 gene and demonstrate fusion of one or multiple calvarial sutures.^25^ While skull development is affected in these syndromes, changes in cartilage, brain, upper face, airway, and mandible morphology have also been noted. ^25–31^ Thus, the impact of these mutations is widespread and could involve changes to other factors within the sutural niche, such as nerves and vasculature.

Imaging the spatial distribution of nerves provides insights into how their interactions with other cell types change with age or disease. Recently, confocal imaging in combination with thick cryosection immunostaining was used to generate a neuroskeletal atlas of the murine long bone periosteum, which identified three patterns of periosteal innervation.^32^ In contrast, there remains a knowledge gap in our understanding of the complete 3D distribution of nerves in the skull, how they change with time, and their association with the vasculature. Similar to long bone, early studies in the murine calvarium identified nerves in small regions of the periosteum using traditional histology.^16,21^ Similarly, histologic cross-sections have been used to show that nerves traverse throughout the interparietal, parietal, and frontal bones, with nerves in the periosteum and dura mater that branched into the diploë regions. However, due to limitations in 3D imaging of large sections of tissue at high spatial resolution^33^, a complete quantitative characterization of nerve distribution in the calvaria has not been feasible until now. A recently developed quantitative lightsheet microscopy (QLSM) workflow provided high-resolution 3D images of the entire murine calvarium.^34^ Analyzing the resulting images provides extensive quantitative data on the distribution and interactions of fluorescently stained cells or anatomic structures. Here, we used an analogous QLSM workflow to map the 3D architecture of nerves (TUBB3+, NeuF+, CGRP+, and TH+) and their spatial associations with vascular phenotypes over the entire murine calvarium over the primary lifespan (i.e. P0-80wk) of C57BL/6J mice.

We found a significant increase in nerve density from P0 to early adulthood (12wk) and subsequent decreases in nerve density as the mice aged, thereby demonstrating for the first time the impact of aging on calvarial nerve. We identified elevations in unmyelinated sensory (CGRP+) and sympathetic (TH+) nerve densities during skeletal maturation of perinatal mice. For all the nerve markers analyzed, we identified their preferential spatial association with Type H blood vessels. Finally, the application of QLSM to a murine model of Apert syndrome, a disease characterized by premature suture closure and skull dysmorphogenesis, revealed that the neurovascular distribution in the calvaria was mildly dysregulated. We believe that these rich 3D imaging data will help to further elucidate the role of calvarial nerves in homeostasis and aging, and aid our understanding of the changes in nerve patterning that occur following bone injury.

## Results

We successfully visualized the 3D distribution of beta-3 tubulin+ (TUBB3+)-stained nerves in the parietal and frontal bones and in the coronal and sagittal sutures using QLSM **(Figure 1A)**. Tissue clearing without decalcification (using 2’2-thiodiethanol) was effective for all murine ages studied and enabled effective visualization of 3D nerve architecture throughout the thickness of the calvarium **(Supplemental Figure 1A)**. Due to the age-dependent background optical signal in the murine calvaria, we developed a pipeline based on the Ilastik®^35^ machine learning software to segment nerve structures with subsequent image analysis with Imaris® **(Supplemental Figure 1B)**. Briefly, this pipeline followed our previous method of image conversion and stitching using Imaris® software,^34^ followed by cropping of the parietal bone, coronal suture, and frontal bone regions for region-specific analyses. Following export of the images into the ImageJ software^36^ and color channel separation, z-stacks were segmented using Ilastik®, which resulted in more accurate segmentation of nerve structure. These segmented nerve data were then imported back into Imaris® for downstream volume and spatial analyses. When the Ilastik® pipeline was compared to that from Imaris®, we observed that nerve structures were more clearly visible, background noise was reduced, and quantitative data was accurate **(Supplemental Figure 1C-D)**. Thus, this pipeline was combined with our blood vessel workflow^34^ for all subsequent analyses.

**Figure 1:**
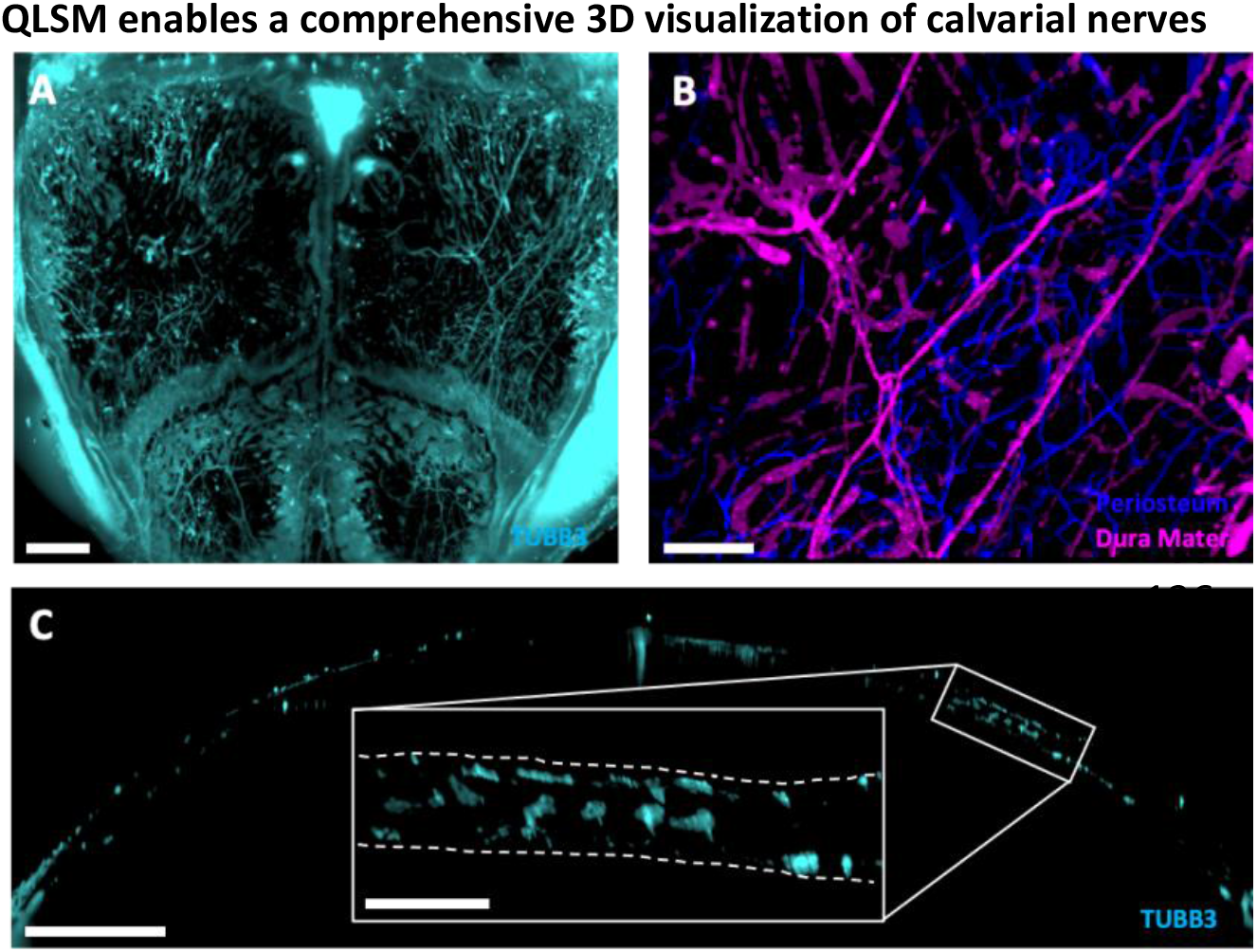
Quantitative lightsheet microscopy (QLSM) enabled 3D visualization of cranial nerves. A) Maximum intensity projection (MIP) image of TUBB3+ nerves in the entire murine calvarium. Scale bar is 1500 μm. B) MIP image of the distribution of parietal nerves in the periosteum (blue) and dura mater (magenta). Scale bar is 300 μm. C) 50 μm thick coronal cross-section of the calvarium acquired with QLSM illustrating TUBB3+ nerves. Scale bar is 1000 μm. Inset scale bar is 300 μm.

In addition to using Imaris® to compute volume and spatial association data, we also used its surface segmentation classification feature to identify nerves that were localized to either the periosteum or the dura mater **(Figure 1B, Supplemental Video 1)**. Next, we explored quantitative differences in nerves within these subregions and explored the changes in their localization to these regions during aging. In comparison to previous methods for visualizing calvarial nerves, our method enabled the 3D visualization of nerves throughout the entire murine calvarium. Although thick cryosections enable adequate visualization of nerves in 2D within specific regions, their 3D structures or interactions can be altered during sectioning. Using QLSM, we could generate cross-sectional images similar to those from thick cryosections, while also generating a 3D image of nerve distribution throughout the frontal and parietal bones without decalcification, dehydration, or geometric distortion **(Figure 1C, Supplemental Figure 2A)**.

To evaluate the ability of the imaging workflow to visualize the network of nerves in the skull at high resolution, we also used confocal imaging to visualize TUBB3+ nerves at both 10× and 20× magnification **(Supplemental Figure 2B-C)**. At 10× magnification (i.e. 1.25 μm resolution), which is similar to the resolution we employed for QLSM (1.3 μm resolution), nerves exhibited similar patterns in each of the three regions **(Supplemental Figure 2B)**. At 20× magnification (i.e. 0.6 μm resolution), we used the DiameterJ plugin in ImageJ® and determined the average nerve diameter to be 4.01±1.57 μm (n = 3) **(Figure 2C)**, which was in agreement with previous studies in sensory and sympathetic nerves.^37,38^ Assuming a normal nerve diameter distribution, this suggests that QLSM captures nearly 95% of the TUBB3+ nerves.

**Figure 2:**
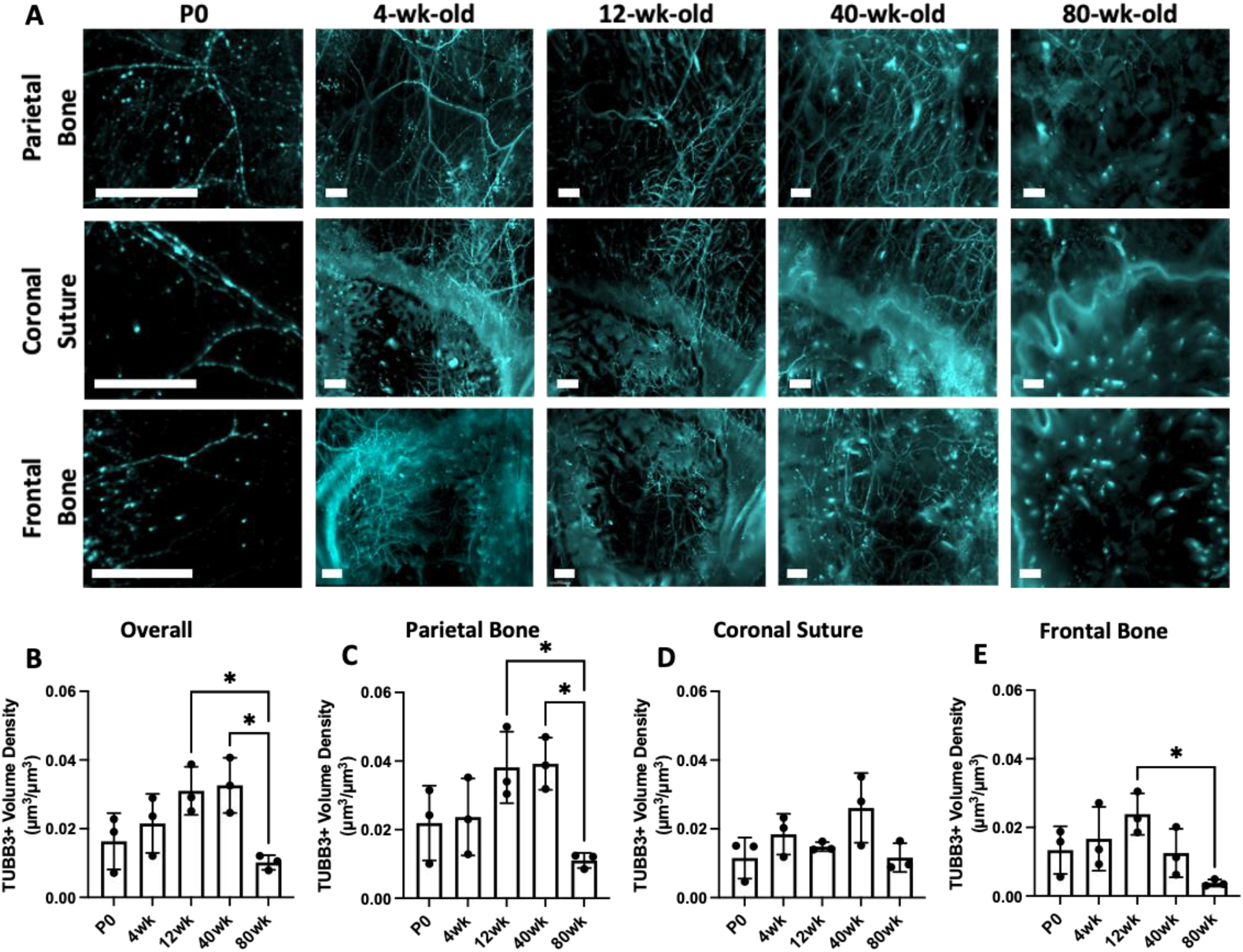
Changes in TUBB3+ nerve distribution over the mouse lifespan. A) MIP images of the parietal bone, coronal suture, and frontal bone regions of P0, 4wk, 12wk, 40wk, and 80wk mice with TUBB3+ nerves. Scale bar is 300 μm. Volume fraction calculation for P0, 4wk, 12wk, 40wk, and 80wk TUBB3+ nerves in B) Overall calvaria, C) Parietal Bone Region, D) Coronal Suture Region, and E) Frontal Bone Region, respectively. Data are mean ± SD. Differences were detected using a two-way ANOVA with post-hoc Tukey HSD (Honestly Significant Difference) test. *p<0.5 where designated.

### Calvarial nerve density and spatial distribution varies across murine lifespan

Using TUBB3 labeling, we imaged the distribution of calvarial nerves at post-natal day 0 (P0), 4wk, 12wk, 40wk, and 80wk calvaria to represent neonatal, developing, skeletally mature, middle-aged, and aged timepoints, respectively. From week 4 onwards, the parietal bone regions exhibited a mesh-like network throughout the bone **(Figure 2A)**. The coronal suture region exhibited nerves that appeared to originate in the suture or along the borders of the bone and projected into the frontal and parietal bones. In aged samples, few nerves were found in the coronal suture region with one large nerve traversing the suture itself. At all ages, the frontal bone exhibited nerves that originated from nearby sutures and traveled radially into the center of the frontal bone **(Figure 2A)**. While we did not quantitatively analyze nerves in the sagittal and interfrontal sutures, we also observed nerves that originated from these sutures **(Supplemental Figure 3D)**. For example, in the sagittal suture, a large nerve bundle was wrapped around the sagittal sinus and expressed high levels of TUBB3 staining at all ages.

**Figure 3:**
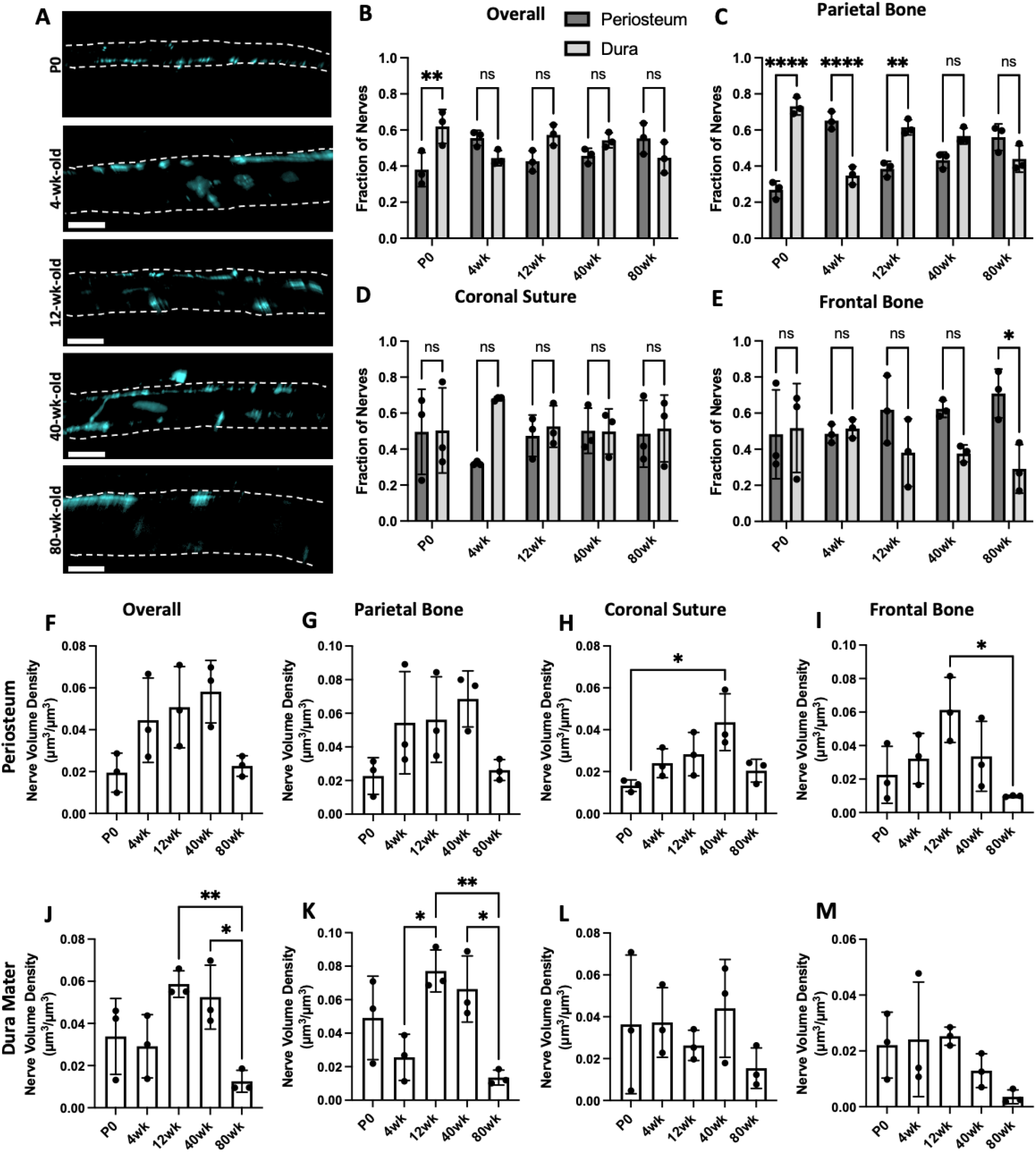
Spatial changes in periosteum and dura mater TUBB3+ nerve distribution over the mouse lifespan. A) 50 μm coronal cross-sections of TUBB3+ nerves in P0, 4wk, 12wk, 40wk, and 80wk mice. Scale bar is 300 μm. Nerve fraction calculations for TUBB3+ periosteal and dural nerves at P0, 4wk, 12wk, 40wk, and 80wk in B) Overall Calvaria, C) Parietal Bone, D) Coronal Suture, and E) Frontal Bone. TUBB3+ periosteal nerve volume density calculations at P0, 4wk, 12wk, 40wk, and 80wk in F) Overall Calvaria Periosteum, G) Parietal Bone Periosteum, H) Coronal Suture Periosteum, and I) Frontal Bone Periosteum. TUBB3+ dural nerve volume density calculations at P0, 4wk, 12wk, 40wk, and 80wk in J) Overall Calvaria Dura Mater, K) Parietal Bone Dura Mater, L) Coronal Suture Dura Mater, M) Frontal Bone Dura Mater. Data are mean ± SD. Statistics were performed with a two-way ANOVA with post-hoc Tukey HSD test. *p<0.5, **p<0.1, ****p<0.001 where designated.

Following quantitative analysis, no difference in nerve volume density was seen between P0, 4wk, 12wk, or 40wk samples, while there was a steep drop-off in nerve volume density in 80wk samples **(Figure 2A)**. This trend was consistent in the parietal bone, but not in the coronal suture or frontal bone **(Figure 2C-E)**. The coronal suture did not demonstrate any significant differences in overall nerve volume density with age **(Figure 2D)**. In the frontal bone, there was a significant decrease in nerve volume density from 12wk to 80wk, but there was also a trend towards a decrease in 40wk samples, as nerve volume density was not maintained throughout 40wk **(Figure 2E)**. This may be indicative that nerve loss with aging may occur faster in the frontal bone than in the parietal bone.

In cross-sectional images, we visualized nerves traveling through transcortical canals with and without blood vessels **(Supplemental Figure 3C, yellow and white arrowheads)**. Using Imaris’® machine learning classification feature, we quantified periosteal and dural nerve populations at all ages. Overall, only P0 samples demonstrated a significant difference between periosteal and dural nerves proportions **(Figure 3A-B)**. The lack of differences at the other ages may be due to differing distributions between bone regions or between nerve subtypes. For example, in 4wk samples, there was a significantly higher proportion of nerves in the periosteum relative to the dura mater of the parietal bone regions, while the converse was true for the coronal suture region **(Figure 2C-E)**. When comparing periosteal and dural volume densities between ages, we found an overall and parietal bone age-dependent increase in the periosteum through 40wk, with a steep drop off at 80wk **(Figure 3F-G)**. In comparison, in the dura mater, there was an increase in dural nerves from 12wk and 40wk, but it was not consistent throughout aging, as no increase was observed between P0 and 4wk samples **(Figure 3J-K)**. This may be indicative that dural nerves may develop later than periosteal nerves and may continue to develop into adulthood. In the coronal suture region, there was an age-dependent increase in nerve volume density through 40wk in the periosteum, while the dura mater demonstrated no significant trends **(Figure 3H, L)**. The lack of differences in the dura mater may explain the lack of differences observes in the coronal suture overall, even though the periosteum exhibited age-dependent changes **(Figure 2D)**. In the frontal bone, in both the periosteum and dura mater, there is an age-dependent decrease in nerve volume density from 12wk samples to 80wk samples, suggesting that nerves regress in the frontal bone periosteum and the dura mater with aging **(Figure 3I-M)**.

### Nerves preferentially associate with shifting CD31^hi^Emcn^hi^ vessels populations over the murine lifespan

We have previously mapped CD31^hi^Emcn^hi^ blood vessels, along with CD31^hi^Emcn^-^ and CD31^lo^Emcn^hi^ blood vessels in the young and adult murine calvaria.^34^ Further, we identified the spatial association between CD31^hi^Emcn^hi^ vessels and osteoprogenitors. In this study, we hypothesized that nerves may demonstrate a preferential association with CD31^hi^Emcn^hi^ vessels due to their known roles in bone development and healing. To first investigate the blood vessel phenotype densities with aging, P0, 4wk, 12wk, 40wk, and 80wk calvaria were stained with CD31 and Emcn **(Figure 4A, Supplemental Figure 3C)**. CD31+ vessels were found throughout the calvaria with similar morphologies at all ages. CD31+ vessels increased in diameter and slightly decreased in volume density with aging, although not significantly **(Figure 4A, Supplemental Figure 3A, D)**. Emcn+ vessels demonstrated drastic morphological changes with aging, as bone marrow-resident sinusoidal blood vessel signal increases with age, while the Emcn^hi^ signal virtually disappears **(Figure 4A, Supplemental Figure 3A, D)**. When quantified, the Emcn^hi^ vessel density increased until skeletal maturation at 12wk, and then declined as aging progressed. Due to the poor signal-to-noise ratio (SNR) of Emcn^lo^ vessels, we did not quantify the Type L (CD31^lo^Emcn^lo^) phenotype.

**Figure 4:**
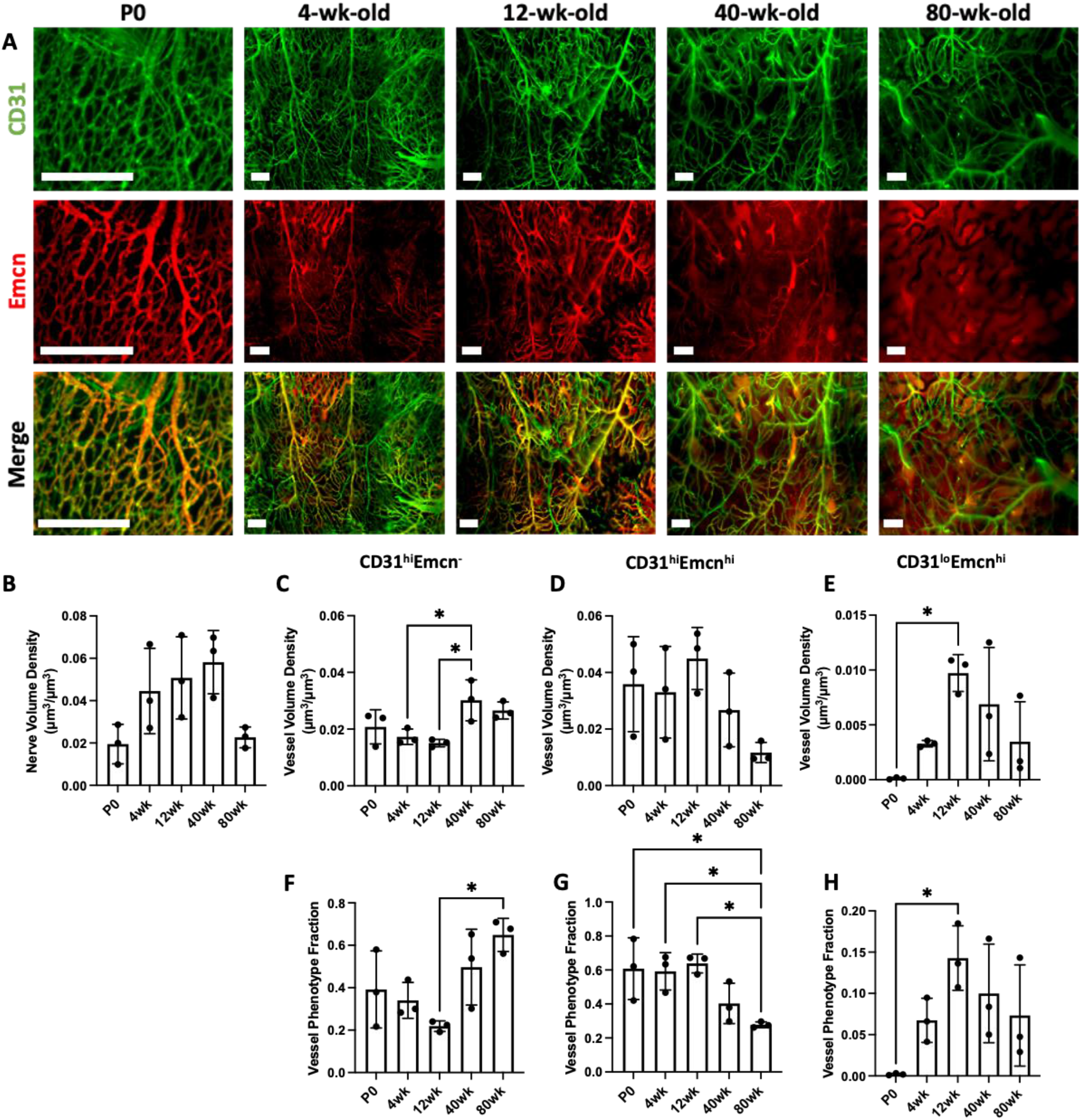
Changes in the spatial distribution of vascular phenotypes during aging. A) MIP images of the parietal bone region of P0, 4wk, 12wk, and 80wk calvaria with vasculature stained with CD31 (green) and Emcn (red). Scale bar is 300 μm. B) Vessel volume density calculation for P0, 4wk, 12wk, 40wk, and 80wk mice. C-E) Phenotypic vessel volume density calculation for C) CD31^hi^Emcn^-^, D) CD31^hi^Emcn^hi^, and E) CD31^lo^Emcn^hi^ vessels in P0, 4wk, 12wk, 40wk, and 80wk mice. F-H) Phenotypic vessel phenotype fraction calculation for F) CD31^hi^Emcn^-^, G) CD31^hi^Emcn^hi^, and H) CD31^lo^Emcn^hi^ vessels in P0, 4wk, 12wk, 40wk, and 80wk mice. Data are mean ± SD. Statistics were performed with a two-way ANOVA with post-hoc Tukey HSD test. *p<0.5 where designated.

Overall blood vessel density remained unchanged with aging **(Figure 4B)**. However, blood vessel phenotypes demonstrated different spatial patterns with aging **(Figure 4C-H, Supplemental Figure 4)**. CD31^hi^Emcn^-^ vessels increased in density and phenotype fraction with aging **(Figure 4C, F)**, while CD31^hi^Emcn^hi^ vessels significantly decreased phenotype fraction me **(Figure 4D, G)**. Although CD31^lo^Emcn^hi^ vessels were found to increase in density from P0 to 12wk **(Figure 4E, H)**, their volume was substantially smaller than for all other phenotypes. This may imply the need for additional samples to elucidate differences between conditions. To ensure an unbiased estimate of blood vessel phenotypes, high and low measurements for CD31 and Emcn blood vessel phenotypes were generated using binary masks. Following masking, CD31^hi^Emcn^-^ vessels appeared green, CD31^hi^Emcn^hi^ vessels yellow, and CD31^lo^Emcn^hi^ vessels red **(Supplemental Figure 5)**. Overall vessel volume and vessel phenotype volumes were then generated from the spatially down-sampled segmented images.

To visualize the spatial relationship of nerves with blood vessel phenotypes, CD31+ and Emcn^hi^ blood vessel staining was combined with TUBB3+ nerve staining (**Figure 5A)**. While some nerves were seen traveling adjacent to larger vessels, most nerves did not directly track alongside vessels in the calvaria **(Figure 5A)**. Overall, we observed that the network of nerves resided in the same plane as the blood vessel network but did not exhibit the same spatial patterns. For spatial association analysis, blood vessels and nerves were separated into 10 μm volumes using the Imaris® down-sampled segmented images to calculate the Euclidean distance for each individual section of the nerve segmentation to that of the blood vessels **(Supplemental Figure 5)**.

**Figure 5:**
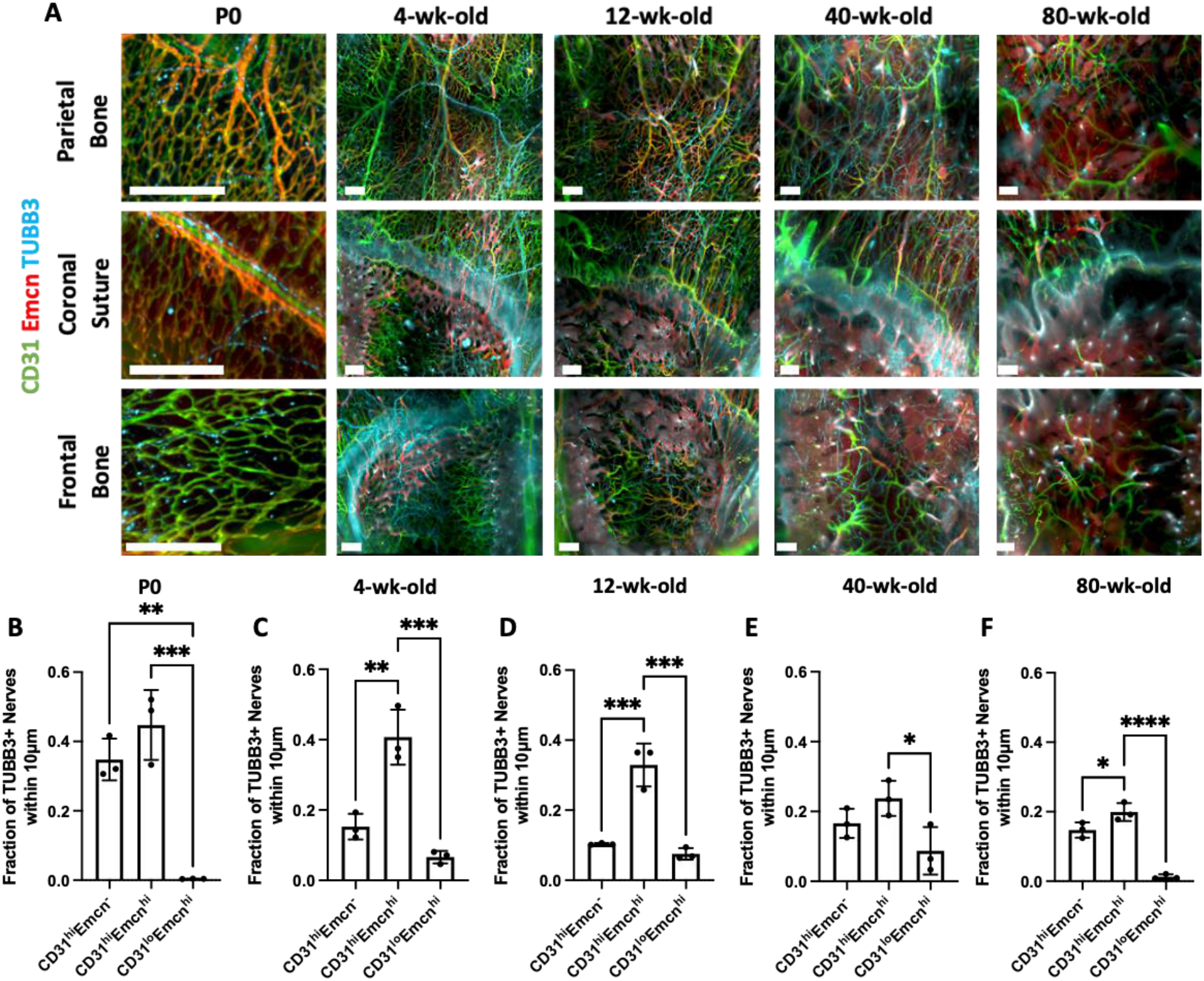
Spatial correlation between vessel phenotypes and TUBB3+ nerves. A) MIP images of the parietal bone, coronal suture, and frontal bone regions of P0, 4wk, 12wk, 40wk, and 80wk mice with TUBB3+ nerves (blue) and vasculature stained with CD31 (green) and Emcn (red). Scale bar is 300 μm. B) The fraction of nerves within 10 μm of each vessel phenotype is displayed for B) P0, C) 4wk, D) 12wk, E) 40wk, and F) 80wk. Data are mean ± SD. Statistics were performed with a two-way ANOVA with post-hoc Tukey HSD test. *p<0.5, **p<0.1, ***p<0.01, ****p<0.001 where designated.

Following classification of vessel phenotypes, nerves were found to spatially associate with CD31^hi^Emcn^hi^ vessels for all ages studied, although the degree of association significantly reduced with aging **(Figure 5B-F, Supplemental Figure 3E-I)**. At all adult ages other than 40wk, this trend was significantly higher than for CD31^hi^Emcn^-^ vessels. This trend was not exclusively due to the prevalence of each vessel phenotype, as CD31^hi^Emcn^-^ vessels were equally or even more prevalent than CD31^hi^Emcn^hi^ vessels in 40wk and 80wk calvaria, respectively **(Figure 4C-D)**. In P0 samples, TUBB3+ nerves were also associated with CD31^hi^Emcn^-^ vessels to the same extent as 4wk and 12wk ages. These data imply that both vessel phenotypes may be communicating with TUBB3+ nerves at neonatal stages.

### Small, unmyelinated nerve subtypes increase with postnatal development, while large, myelinated nerves remain unaffected

While TUBB3+ nerves were impacted by aging, we did not find any substantial differences in TUBB3+ nerve distributions or spatial distribution during skeletal maturation that occurs in early adulthood. Murine bones continue to develop after birth and are considered skeletally mature around 10 weeks of age.^39^ Furthermore, previous studies have determined that nerves are essential to proper bone development.^1^ Thus, we explored changes in sensory markers, neurofilament (NeuF) and calcitonin gene-related peptide (CGRP), and sympathetic marker, tyrosine hydroxylase (TH), during (i.e. 4wk) and after (i.e. 12wk) postnatal development and skeletal maturation. CGRP+ and TH+ nerves demonstrated a higher density following postnatal development, while NeuF+ nerves were unchanged, suggesting that some nerve subtypes may expand, mature, and specialize along with bone maturation **(Figure 6A-B, F-G, K-L, Supplemental Figure 6)**. Overall, increases in CGRP+ and TH+ nerves seemed to be consistent throughout the calvaria with slight increases found in the parietal bone, coronal suture, and frontal bone for both markers **(Supplemental Figure 5)**.

**Figure 6:**
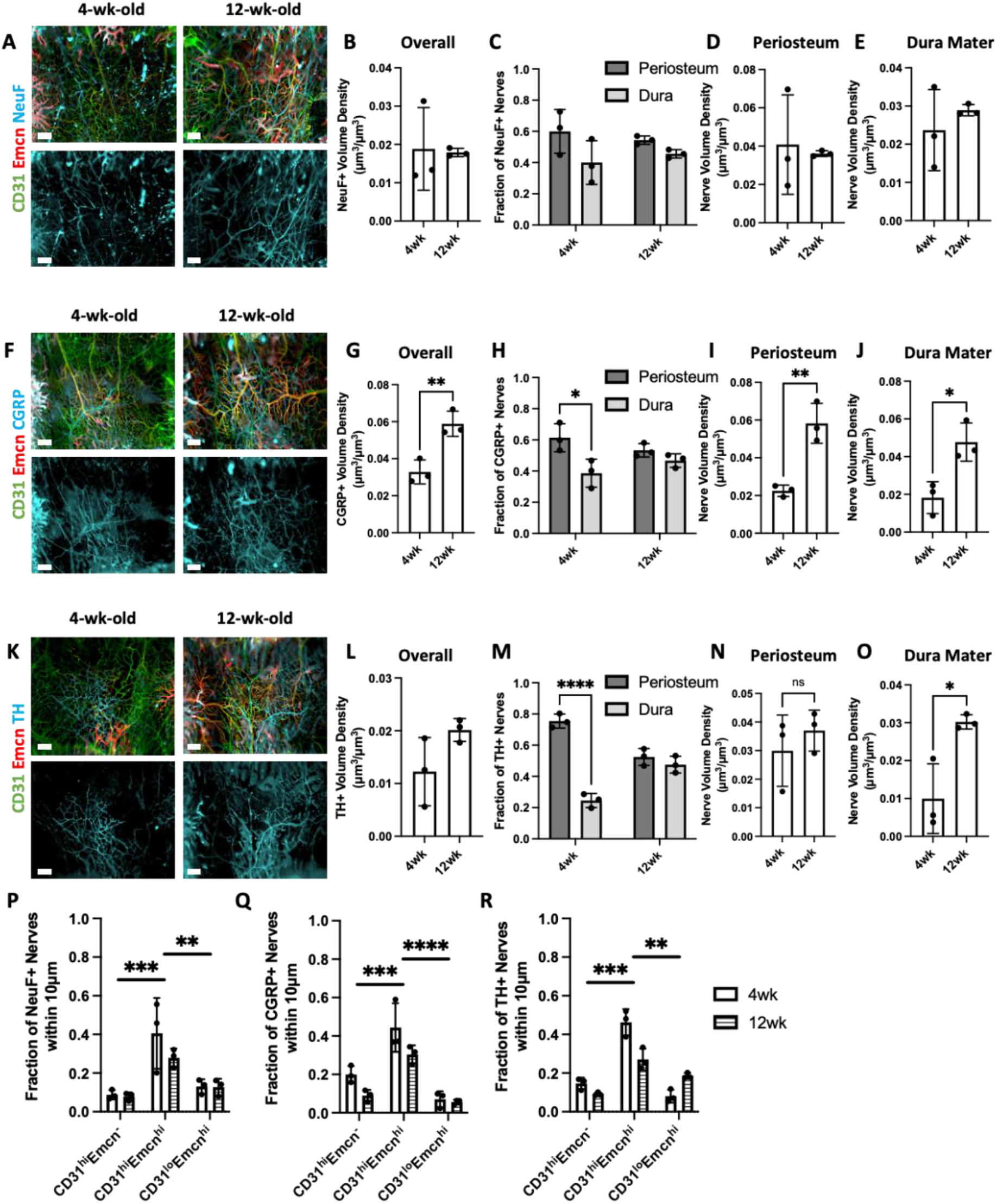
Changes in nerve subtype distribution and neurovascular interactions during postnatal development. A) MIP images of the parietal bone region of 4wk and 12wk mice with NeuF+ nerves (blue) and vasculature stained with CD31 (green) and Emcn (red). Scale bar is 300 μm. B) Nerve volume fraction calculation for 4wk and 12wk NeuF+ nerves. C) Periosteal and dural nerve fraction calculations for NeuF+ nerves in 4wk and 12wk mice. D-E) Periosteal and dural nerve volume density calculations for NeuF+ nerves in 4wk and 12wk samples in D) Overall Calvaria Periosteum, E) Overall Calvaria Dura Mater. F) Maximum intensity projections of the parietal bone region of 4wk and 12wk mice with CGRP+ nerves (blue) and vasculature stained with CD31 (green) and Emcn (red). Scale bar is 300 μm. G) Nerve volume fraction calculation for 4W and 12wk CGRP+ nerves. H) Periosteal and dural nerve fraction calculations for CGRP+ nerves in 4wk and 12wk mice. I-J) Periosteal and dural nerve volume density calculations for CGRP+ nerves in 4wk and 12wk samples in I) Overall Calvaria Periosteum, J) Overall Calvaria Dura Mater. K) Maximum intensity projections of the parietal bone region of 4wk and 12wk mice with TH+ nerves (blue) and vasculature stained with CD31 (green) and Emcn (red). Scale bar is 300 μm. L) Nerve volume fraction calculation for 4wk and 12wk TH+ nerves. M) Periosteal and dural nerve fraction calculations for TH+ nerves at 4wk and 12wk. N-O) Periosteal and dural nerve volume density calculations for TH+ nerves in 4wk and 12wk samples in N) Overall Calvaria Periosteum, O) Overall Calvaria Dura Mater. P-R) Spatial association calculations for nerves and vessels phenotypes for P) NeuF+, Q) CGRP+, and R) TH+ nerves. The fraction of nerves within 10 μm is displayed for each vessel phenotype. Data are mean ± SD. Statistics were performed with a two-way ANOVA with post-hoc Tukey HSD test and a two-tailed t-test. *p<0.5, **p<0.1, ***p<0.01, ****p<0.001 where designated.

In terms of their distribution in the periosteum and dura mater, CGRP+ and TH+ nerves demonstrated a preferential localization to the periosteum at 4wk; however, this localization was lost after postnatal development in both cases **(Figure 6H, M, Supplemental Figure 7)**. Interestingly, this trend was consistent in the parietal bone for both CGRP+ and TH+ nerves and in the coronal suture for TH+ nerves, but no changes were seen in the other regions **(Supplemental Figure 7)**. When periosteum and dura mater nerve volumes were considered, we found an increase in nerve volume density in both the periosteum and dura mater for CGRP+ nerves, but only an increase in the dura mater for TH+ nerves **(Figure 6D-E, I-J)**. This trend was consistent throughout the regions of the calvaria. CGRP+ nerves increased in every region with postnatal development, while TH+ nerves exhibited only regional increases in the dura mater **(Supplemental Figure 8)**. Thus, the unmyelinated nerve growth after postnatal development preferentially occurred in the dura mater for TH+ nerves and throughout the periosteum and dura mater for CGRP+ nerves. No differences were observed for dura and periosteum distributions of NeuF+ nerves **(Figure 6C)**. For these nerve subtypes, their spatial associations with blood vessel phenotypes were virtually unchanged with postnatal development. NeuF+, CGRP+, and TH+ nerves all maintained their association with CD31^hi^Emcn^hi^ vessels within 10 μm **(Figure 6P-R)**.

### The Fgfr2 P253R mutation associated with Apert syndrome demonstrates some impact on neurovascular architecture

Nealy 100% of all cases of Apert syndrome are caused by one of two FGFR2 mutations and result in coronal suture fusion prior to birth. The *Fgfr2*^*+/P253R*^ *A*pert syndrome mouse is characterized by early mortality with variations in syndrome prevalence. Mice with more moderate phenotypes tend to live up to three weeks, while mice with severe phenotypes live up to only a few days. Thus, for this study, we explored the impact of Apert syndrome on neonatal mice as compared to wild-type (WT) non-mutant litter mates. Calvaria of P0 *Fgfr2*^*+/P253R*^ Apert syndrome mice and their unaffected littermates (*Fgfr2*^*+/+*^) were stained with TUBB3, CD31, and Emcn and imaged with our QLSM workflow. Qualitatively, the neurovascular architecture was maintained between *Fgfr2*^*+/+*^ and *Fgfr2*^*+/P253R*^ samples **(Figure 7A)**. A network of vessels was identified throughout for both groups and nerves were clearly identifiable in all regions. Specifically, in the coronal suture, nerves were found traversing the suture region in both groups. When quantified, there were no discernable differences in nerve distributions between *Fgfr2*^*+/+*^ and *Fgfr2*^*+/P253R*^ groups **(Figure 7B-D)**. Overall and regional nerve volumes remained unchanged **(Figure 7B-C)**, while periosteum and dura mater distribution were mildly impacted. While not significant, there was a higher proportion of nerves in the dura mater than the periosteum of *Fgfr2*^*+/+*^ mice. However, this proportion was lost in *Fgfr2*^*+/P253R*^ samples **(Figure 7D)**. When the volume density of periosteal nerves and dural nerves was compared between groups, no difference was seen between periosteal volumes, while a non-significant decrease was seen in the dura mater **(Figure 7E-F)**. Thus, Apert syndrome may result in nerve loss in the dura mater region following suture fusion.

**Figure 7:**
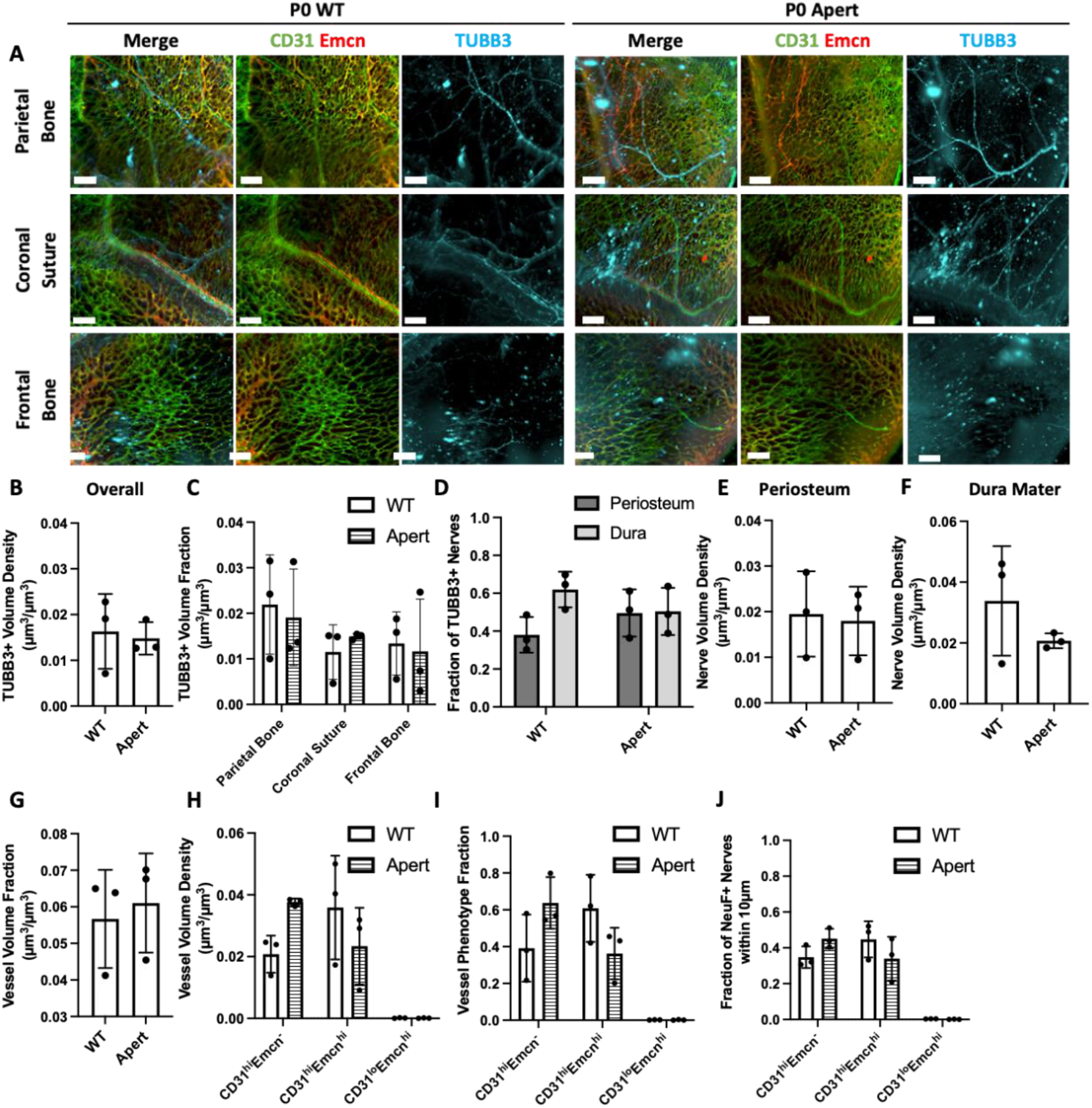
Changes in neurovascular distributions with Apert’s syndrome. A) MIP images of the parietal bone, coronal suture, and frontal bone regions of P0 WT and Apert mice with TUBB3+ nerves (blue) and vasculature stained with CD31 (green) and Emcn (red). B) Overall nerve volume fraction calculation for WT and Apert TUBB3+ nerves. C) Regional nerve volume fraction calculations in the parietal bone, coronal suture, and frontal bone regions for WT and Apert mice. D) Periosteal and dural nerve fraction calculations for TUBB3+ nerves in WT and Apert mice. E-F) Periosteal and dural nerve volume density calculations for TUBB3+ nerves in WT and Apert samples in E) Overall Calvaria Periosteum, F) Overall Calvaria Dura Mater. G) Total vessel volume fraction calculation for WT and Apert mice. H) Phenotypic vessel volume density calculation for CD31^hi^ Emcn^-^, CD31^hi^ Emcn^hi^, and CD31^lo^ Emcn^hi^ vessels in WT and Apert mice. I) Phenotypic vessel phenotype fraction calculation for CD31^hi^ Emcn^-^, CD31^hi^ Emcn^hi^, and CD31^lo^ Emcn^hi^ vessels in WT and Apert mice. J) Spatial association calculations for TUBB3+ nerves and vessels phenotypes. The fraction of nerves within 10 μm is displayed for each vessel phenotype.

Similarly, the vasculature of *Fgfr2*^*+/P253R*^ Apert syndrome mice was only mildly impacted. Overall vessel volumes were unchanged between groups **(Figure 7G)**, while no significant differences were observed between vessel phenotypes. We identified an increase in CD31^hi^Emcn^-^ vessels and a decrease in CD31^hi^Emcn^hi^ vessels **(Figure 7H-I)**. These changes may contribute to the degree of neurovascular association. In *Fgfr2*^*+/P253R*^ Apert syndrome samples, we found a slight increase in association with CD31^hi^Emcn^-^ vessels and a slight decrease with CD31^hi^Emcn^hi^ vessels **(Figure 7J)**. Furthermore, the association with CD31^hi^Emcn^-^ vessels was found at a greater fraction than CD31^hi^Emcn^hi^ vessels in *Fgfr2*^*+/P253R*^ Apert syndrome mice, suggesting that nerves impacted by the Fgfr2 P253R mutation may not be communicating with Type H vessels. Thus, dysregulation of the suture microenvironment may result in changes to the microvasculature, which could impact bone development, homeostasis, and healing capacity.

## Discussion

Here, we present the first quantitative assessment of nerve subtype distributions and their association with vascular phenotypes in the calvaria during postnatal development. Compared to previous studies that explored neonatal and adult calvaria,^21,33^ we demonstrated distinct association of sensory and sympathetic nerves with blood vessels in the skulls of 4wk and 12wk mice. Other than in the transcortical canals, we did not observe nerves in cortical bone. Following postnatal development, CGRP+ and TH+ nerve volume fractions increased with skeletal maturation, suggesting that nerves may further specialize as the skeleton matures. This increase in CGRP+ and TH+ nerves was similar to the increase in the percentage of CGRP+ nerves found in the dorsal root ganglia (DRG) from P12 to adulthood.^40^ CGRP is known to stimulate osteoblast proliferation and differentiation, while inhibiting osteoclast formation.^41,42^ Thus, CGRP+ nerves likely play a role in maintaining bone turnover and structure in mature bone. As for TH+ nerves, expression of TH increased in the superior cervical ganglion, olfactory bulb, and brain following postnatal development.^43–45^ The increase in TH+ expression was associated with tissue maturation and was likely similar in the calvarium. Further studies will be required to address calvarium-specific CGRP+ and TH+ neurons that develop with postnatal development, as they may play significant roles in the response to environmental stressors and bone maintenance. Interestingly, we did not find any differences in NeuF+ nerves with postnatal development. Previous studies in the trigeminal ganglia have suggested that large diameter neurons mature prior to small diameter neurons,^46^ which may explain why the small, unmyelinated CRGP+ and TH+ nerves continue to increase as the skeleton matures, while large, myelinated NeuF+ nerves do not.

Age-related changes in nerves have been characterized in long bones. Chartier et al. reported a decrease in the number of nerves in the periosteum of murine femurs, but no decrease in nerve density, due to the thinning of the periosteum with age.^47^ Using multivariate analysis, Steverink et al. (2021) found an age-related decrease in nerves in human bones that was most significant in the periosteum.^10^ Despite differences in embryonic origin and developmental pathways of calvarial and post-cranial bone, the calvaria also exhibited decreased nerve density with aging, specifically in parietal and frontal bone regions, but not in the coronal suture region. Due to the ability of QLSM to provide ‘global’ 3D readouts of the murine calvaria, we could investigate differences throughout the full calvaria and quantitatively characterize specific regions. Thus, we obtained unique insight into age-related nerve loss in the calvaria, which occurred earlier in the frontal bone compared to the parietal bone. Since nerves have been shown to be necessary for the maintenance of bone homeostasis and skeletal stem cell proliferation,^24,48^ this nerve loss may be associated with age-related bone loss and enhanced bone fragility. In contrast, nerves may also become less necessary for aged bone homeostasis, since Wu et al. (2016) identified a reduced trabecular and cortical thinning after inferior alveolar nerve transection in aged mice as compared to young and middle-aged mice.^49^ Thus, nerves may not mediate homeostasis to the same extent in aged individuals as compared to highly regenerative young or densely innervated middle-aged individuals. Previous studies have identified nerves in both the periosteum^16,33,50–52^ and dura mater,^21,33,50,51^ with one study suggesting that there is a higher density of CGRP staining in the dura mater.^33^ Therefore, we further stratified nerve populations as belonging to the periosteum or dura mater. Conversely, we identified a higher CGRP+ nerve population in the periosteum compared to the dura mater in young mice (4wk). Nevertheless, the spatial distribution of nerves in the periosteum and dura mater varied depending on their location (i.e. parietal bone, coronal suture, and frontal bones). This might explain some of the discrepancies with prior findings.

We also identified preferential spatial association of nerves with CD31^hi^Emcn^hi^ vessels in orthotopic bone which is maintained throughout murine aging. In a previous study by Qin et al., the number of CD31^hi^Emcn^hi^ vessels was reduced following TrkA inhibition during heterotopic ossification, suggesting that neuron-to-vessel signaling may mediate CD31^hi^Emcn^hi^ vessel development in ectopic bone formation and may also be important in orthotopic bone development, homeostasis, and healing. In general, it is well known that nerves control vessel patterning, specifically for arteries and arterioles.^11–13^ Further, NGF/TrkA signaling in sensory neurons resulted in VEGF secretion and angiogenesis.^53,54^ Previous studies have identified a role for TrkA+ nerves in controlling vascularization during development, response to mechanical loading, and bone healing.^1,48,51,55^ To the best of our knowledge, there have been no studies that have explored the importance of vessel phenotype in these interactions. However, as CD31^hi^Emcn^hi^ vessels are also positively associated with bone development and healing,^22^ they are likely the forerunners in neurovascular signaling. Although further studies are necessary to discern the interactions between nerves and CD31^hi^Emcn^hi^ vessels in the calvarium, this study provides strong justification for the importance of these interactions. Interestingly, although CD31^hi^Emcn^hi^ blood vessels were reduced in aged mice, their preferential spatial association was maintained throughout all ages studied, albeit at a lower level, emphasizing the importance of this interaction.

In contrast to our findings, a prior study^56^ was unable to identify a network of blood vessels throughout the full parietal and frontal bones for adult ages (10-14 weeks), whereas we found a dense network of vessels throughout the entire tissue. This discrepancy may be due to their focus on the bone marrow region while excluding the periosteum and dura mater. In fact, we found that CD31^hi^Emcn^-^ blood vessels increased in density and fraction with age, while CD31^hi^Emcn^hi^ blood vessels decreased in fraction with age, which was in agreement with previous studies in long bone.^6^ Loss of CD31^hi^Emcn^hi^ blood vessels is associated with reduced osteogenic-angiogenic coupling and regenerative capacity. Differences in methodology may account for some of the differences in overall blood vessel volume found between our studies. The Adams et al. study segmented both Emcn^hi^ and Emcn^lo^ blood vessels, whereas we exclusively segmented Emcn^hi^ blood vessels and did not take the CD31^lo^Emcn^lo^ phenotype into account due to its low SNR.

In addition to aging, we also investigated the impact of craniofacial disease associated with genetic mutations on neurovascular architecture in a model of Apert syndrome, which demonstrates early coronal suture fusion *in utero*. We identified a non-significant increase in CD31^hi^Emcn^-^ vessel volume and a non-significant decrease in CD31^hi^Emcn^hi^ vessel volume. In a study of Crouzon syndrome, another FGFR-related craniosynostosis syndrome that demonstrates premature suture fusion, an increase in blood vessel diameter was found in the cranial vault of four human patients.^57^ As CD31^hi^Emcn^-^ vessels are generally arteriole-like, since they do not express Emcn, and CD31^hi^Emcn^hi^ vessels are primarily capillary-like, our preclinical results demonstrate the similar findings between two craniosynostosis syndromes. In comparison, in a study of TWIST1 mutations, which also demonstrates craniosynostosis, normal arterial development was identified, suggesting that impacts on the vasculature during craniosynostosis may be mutation specific.^58^ While Apert syndrome patients demonstrate abnormalities in the central nervous system,^59,60^ little is known its impact on peripheral nerves in the calvaria, specifically those in the impacted calvaria. In this study, although we found a non-significant increase in dural nerves with Apert syndrome, we did not identify an overall change to calvarial nerves with Apert syndrome. As this is the first identification of changes to peripheral nerves with Apert syndrome and because suture fusion occurs prior to P0, additional studies with earlier time points are required to investigate the impact of dural nerve loss on early suture fusion and other Apert syndrome-related anomalies.

Given the close spatial association of the skull and brain along with the direct physical connections between the two, understanding nerve distribution and their vascular association within the skull may provide important insights into developmental abnormalities, age-related cognitive disorders, migraines, or simply, regeneration of injuries. Developing methods that impact the neurovascular architecture can be informative in the design of therapies that promote bone healing. In this study, we acquired the 3D architecture and distribution of nerves throughout the parietal and frontal bones of C57BL/6J mice and quantified changes that occurred over their lifespan. Additionally, we characterized vascular changes throughout the lifespan and the spatial associations of calvarial nerves with specific vascular phenotypes. Notably, we identified varying levels of age-related nerve growth or loss in the parietal and frontal bones, as well as in the periosteum and dura mater. Thus, for studies that target neurovascular infiltration following injury or disease induction, it is important to consider the distribution and densities of nerves and blood vessels in each specific bone or region of interest because they could impact study success and expected outcomes.

## Materials and Methods

### Materials

All antibodies and reagents used in this study can be found in **Supplemental Table 1**.

### Animal models

All animal experiments were approved by the Johns Hopkins University Institutional Animal Care and Use Committee (Protocol No. MO21M146). Animals were housed and cared for in Johns Hopkins’ Research Animal Resources central housing facilities. Young (4wk), adult (12wk), and middle aged (40wk) mice used in this study were C57BL/6J (The Jackson Laboratory, Stock No: 000664). Aged (80wk) mice were a generous gift from Dr. Jeremy Walston. Fgfr2+/P253R Apert syndrome mice and their non-mutant littermates were a generous gift from Dr. Joan Richtsmeier and were generated at Icahn School of Medicine at Mount Sinai and the Pennsylvania State University.^25,61^ Newborn mice (P0) were euthanized by inhalation anesthetics and fixed in 4% paraformaldehyde. Gestation time was 19.0 ± 0.5 days. Genotyping of tail DNA by PCR was performed to distinguish mutant from non-mutant littermates.^25,61^ Mouse litters were produced, sacrificed, and processed in compliance with animal welfare guidelines approved by the Icahn School of Medicine at Mount Sinai and the Pennsylvania State University Animal Care and Use Committees.

### Murine calvarial harvest

Mice were anesthetized with ketamine (100 mg/kg) and xylazine (20 mg/kg). Following anesthesia, mice were injected subcutaneously with 200 U of heparin prior to perfusion. An incision was first made at the xiphoid process and additional incisions were made to gain access to the rib cage and heart. Following disruption of the pleural cavity, heparinized saline (10 U/mL in 1X PBS) was perfused via a 20G blunt needle into the heart via the left ventricle. Prior to perfusion, the right atrium was cut open to allow an open circulation. Perfusion was performed at a rate of 10 mL/min. Following perfusion, the skin was removed around the skull and the calvaria was harvested while maintaining the periosteum and the dura mater. Calvaria were fixed in 4% methanol-free paraformaldehyde overnight at 4°C. Following fixation, calvaria were washed three times for 1 hour with PBS at RT.

### Whole-mount immunostaining and optical clearing

To stain and image calvaria via QLSM, we used a previously published protocol.^34^ Briefly, samples were blocked overnight at 4°C in a blocking buffer containing 10% V/V normal donkey serum and wash buffer (0.1 M Tris, 0.15 M NaCl, 0.05% V/V Tween-20, 20% V/V dimethylsulfoxide, pH 7.5). To enhance signal in the AF555 channel, a biotin blocking kit was used. Calvaria were blocked for 8 hours at RT to block endogenous biotin. Primary staining was performed with antibodies for CD31, Emcn, and TUBB3, NeuF, CGRP, or TH for 7 days at 4°C in blocking buffer. Next, samples were stained with secondary antibodies, either fluorophore- or biotin-conjugated, for 7 days at 4°C. Then, a streptavidin conjugate was used for 5 days to amplify Emcn signal in the AF555 channel. All antibodies and conjugates were diluted in blocking buffer. Between each staining incubation, samples were washed five times for at least 1 hour in wash buffer. Following staining and immediately prior to imaging, samples were cleared in a graded series of 2,2-thiodiethanol (TDE in TBS-Tween; 25%, 50%, 75%, 90%, 100% × 2). Clearing steps were performed for a minimum of 2 hours at RT or overnight at 4°C. Samples were stored in 100% TDE prior to imaging.

### Light-sheet imaging

For lightsheet imaging, we used a LaVision Biotec Ultramicroscope II that was aligned for the refractive index of TDE. Calvaria were mounted to a sample mount and immersed in a glass imaging cuvette with 100% TDE. Samples were imaged using three separate tile acquisitions: a 3 × 1 tile with double-sided illumination, a 3 × 2 tile with left-sided illumination, and a 3 × 2 tile with right sided illumination. Tiles were overlapped by 15% to facilitate stitching. The following hardware and settings were used for all scans: ×2.5 zoom with a ×2 dipping cap (×5 magnification, 1.3 μm x–y pixel size), 5.5 Megapixel sCMOS camera, 20 ms exposure time, 0.154 numerical aperture, and 2.5 μm z step size. Channels were imaged using 561, 640, and 785 nm lasers and 620/60, 680/30, and 845/55 filters, respectively. Laser intensities were set for each antibody and held consistent for each scan.

### Image processing and analysis

All image processing and analysis was performed with Imaris® 9.10, Ilastik®,^35^ and ImageJ^36^ software and a Dell Precision 7820 Tower workstation. The workstation was equipped with a dual Intel Xeon Gold 6240 processor, 384 GB DDR4 SDRAM (2666 MHz speed), 512 GB and 1 TB SATA SSDs, NVIDIA Quadro RTX5000 graphics card (16 GB GDDR6 memory), and Windows 10 Pro for Workstations.

#### Imaris-based Image Conversion and Stitching

To process the raw data for Imaris®, we converted LaVision Biotec raw OME-TIFF files to the Imaris® file format (.ims) using the Imaris® File Converter 9.8. Tiles were manually aligned and stitched using the Imaris® Stitcher 9.8, resulting in a final 3D image. Three pre-defined volumes-of-interest (VOI) were positioned for each data set in the parietal bone, coronal suture, and frontal bone regions. VOI dimensions were consistent between data sets. VOI were cropped and exported into ImageJ for channel separation.

#### Ilastik®-based Image Segmentation

We segmented nerves in all images using Ilastik®, a freely available machine-learning based image segmentation tool.^35^ Briefly, once channels were separated in ImageJ, one z-slice was selected from each region and each condition for Ilastik® training. All features were selected for Ilastik® training to account for changes in intensity, shape, and texture. Individual z-slices were trained for nerve and background signals and prediction masks exported for each region. The remaining z-slices for each condition were segmented using batch segmentation and the corresponding prediction masks. Following Ilastik® segmentation, binary segmentation images were generated in ImageJ. Next, z-stacks are imported back into Imaris® for subsequent analysis.

#### Imaris®-based 3D Nerve Analysis

Nerve channels (stained with TUBB3, NeuF, CGRP, or TH) were segmented using absolute intensity surface segmentation and a 10^4^ μm^3^ volume filter to eliminate subcellular-sized segments. Blood vessels (stained with CD31 or Emcn) were segmented using surface segmentation with a 10 μm radius for background subtraction and a 10^4^ μm^3^ volume filter to eliminate subcellular-sized segments. Thresholds were optimized for each stain and kept consistent between sample ages. Volume statistics for nerves and CD31 and Emcn individual stains were exported from this initial segmentation. Volume density measurements were calculated from the nerve or blood vessel volumes and normalized by the region size.

Next, images were down sampled by a factor of two in each dimension. Down sampling allowed further processing of the initial segmentation to identify vascular phenotypes and nerve localization to the periosteum and dura mater. Binary masks for all stains were generated. A second segmentation (referred to as the down-sampled segmentation) was performed on these masks which included the Imaris® “Split Objects” Surfaces function with a 10 μm seeding point diameter. CD31^hi^Emcn^−^ and CD31^hi^Emcn^hi^ vessels were segmented using the CD31 mask and filtered based upon the absence or presence of masked Emcn signal within each object, respectively. CD31^lo^Emcn^hi^ vessels were segmented based upon the Emcn mask and filtered to remove objects co-localized with masked CD31 signal. Vessel volume fraction measurements were calculated from each phenotype vessel volume and normalized by the overall vessel volume in each sample.

To quantify nerve segments in the periosteum and dura mater, the nerve down-sampled segmentation was further classified. VOIs were analyzed individually, and a training set was selected from the points within each VOI. Initial training data was selected, and iterations were made to ensure an accurate classification. Periosteal and dural nerve fractions were calculated from the nerve section count in either the periosteum or dura mater and normalized by the total nerve count in that region. Spatial association statistics were generated for each vessel phenotype by exporting the shortest distance metrics from the nerve segmentation.

### Statistics

GraphPad Prism was used to perform all statistics. Either a two-tailed t-test or a one-way or two-way ANOVA with post-hoc Tukey HSD test was performed. P-values<0.05 were considered significant.

## Acknowledgements

This work was supported by funding from NIDCR (1R01DE027957), Maryland Stem Cell Research Fund (2022-MSCRFV-5782), the NSF GRFP and NCI (5R01CA237597-05, 2R01CA196701-06A1). Lightsheet imaging was performed at JHU’s Integrated Imaging Center. We thank Joan Richtsmeier for comments on the manuscript.

## Author Contributions

A.L.H. and W.L.G conceived the study. A.L.H. performed all experiments, imaging, and image analysis. Y.R., A.N.R, and A.P. helped design the staining and image analysis protocols. A.L.H. wrote the manuscript. All authors reviewed the manuscript and discussed the work.

