## Supplemental Figures for "3D Imaging Reveals Changes in the Neurovascular Architecture of the Murine Calvarium with Aging"

400 N. Broadway
Smith 5023
Baltimore, MD, 21231

### Supplemental Figures

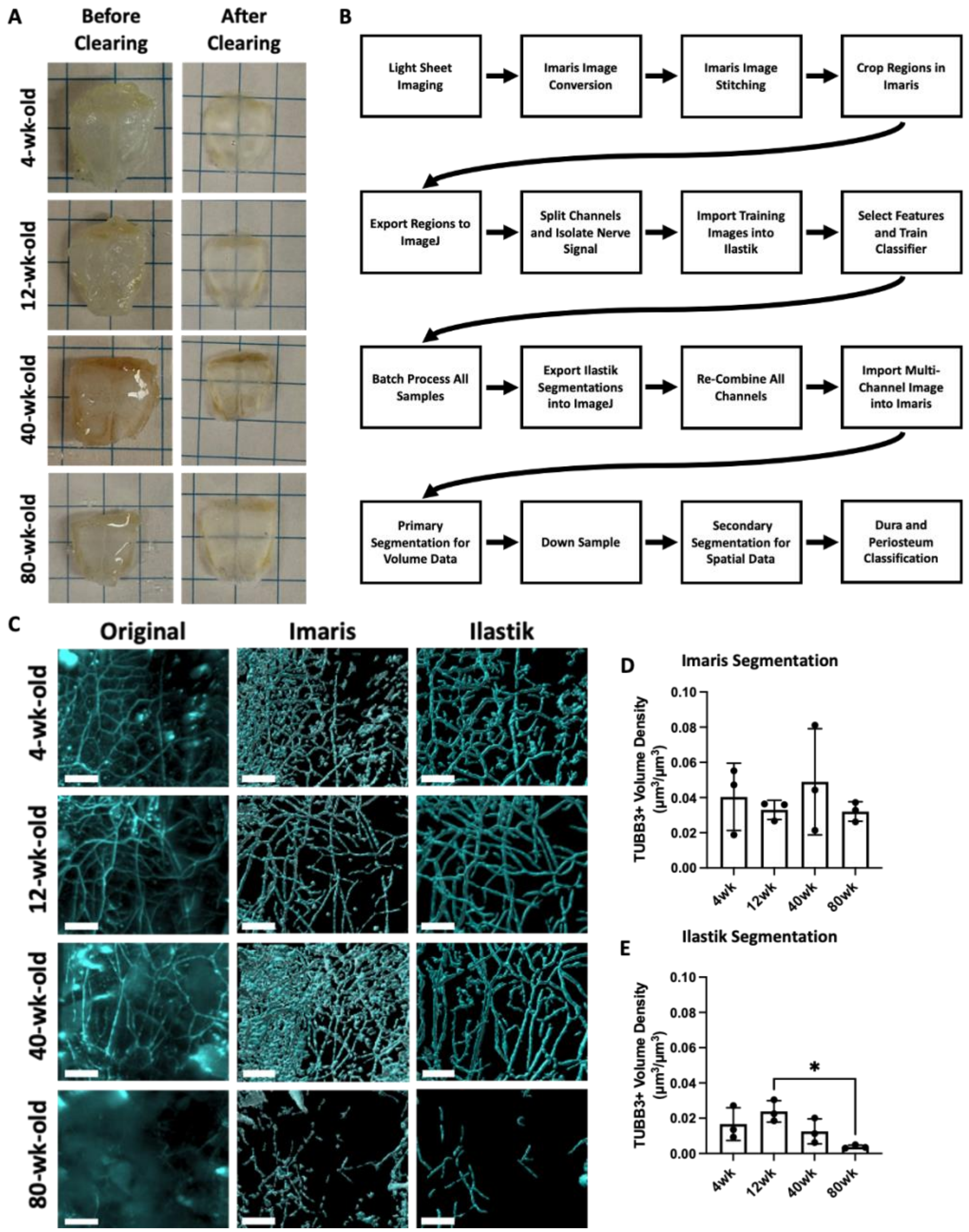


**Supplemental Figure 1: A method for quantitative analysis of 3D nerve structures acquired with QLSM.** A) 4wk, 12wk, 40wk, and 80wk calvaria before and after tissue clearing with 2’2-thiodiethanol. B) Workflow of quantitative nerve analysis beginning with lightsheet imaging, followed image processing with Imaris®, ImageJ®, and Ilastik®. C) Comparison of segmentation of nerves using Imaris® and Ilastik® relative to the original fluorescent images acquired with QLSM. Scale bar is 250 µm. D) Imaris®-derived volume fraction for TUBB3+ nerves in the frontal bone region. E) Ilastik®-derived volume fraction for TUBB3+ nerves in the frontal bone region. Data are mean ± SD. Statistics were performed with a two-way ANOVA with post-hoc Tukey HSD test. *$p<0.5$ where designated.

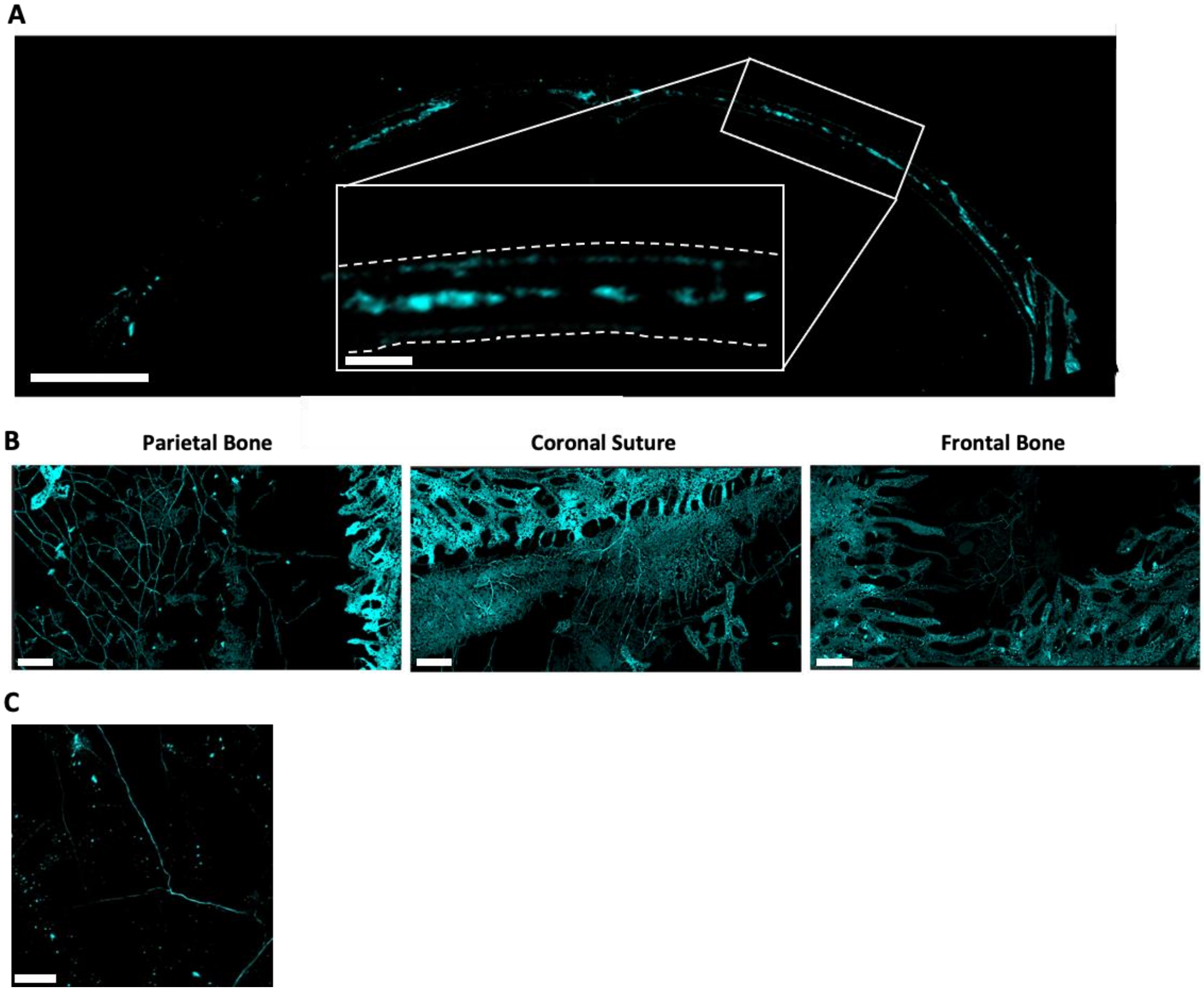


**Supplemental Figure 2: Alternate methods for imaging calvarial nerves.** A) 50 µm thick cross-section of TUBB3+ nerves from a coronal calvarium section. Scale bar is 1000 µm. Inset scale bar is 150 µm. B) MIP image acquired at 10x from confocal microscopy of the whole-mount calvarial tissue in the parietal bone, coronal suture, and frontal bone. Scale bar is 200 µm. C) MIP image acquired at 20× with confocal microscopy of whole-mount calvaria; tissue in the parietal bone. Scale bar is 50 µm.

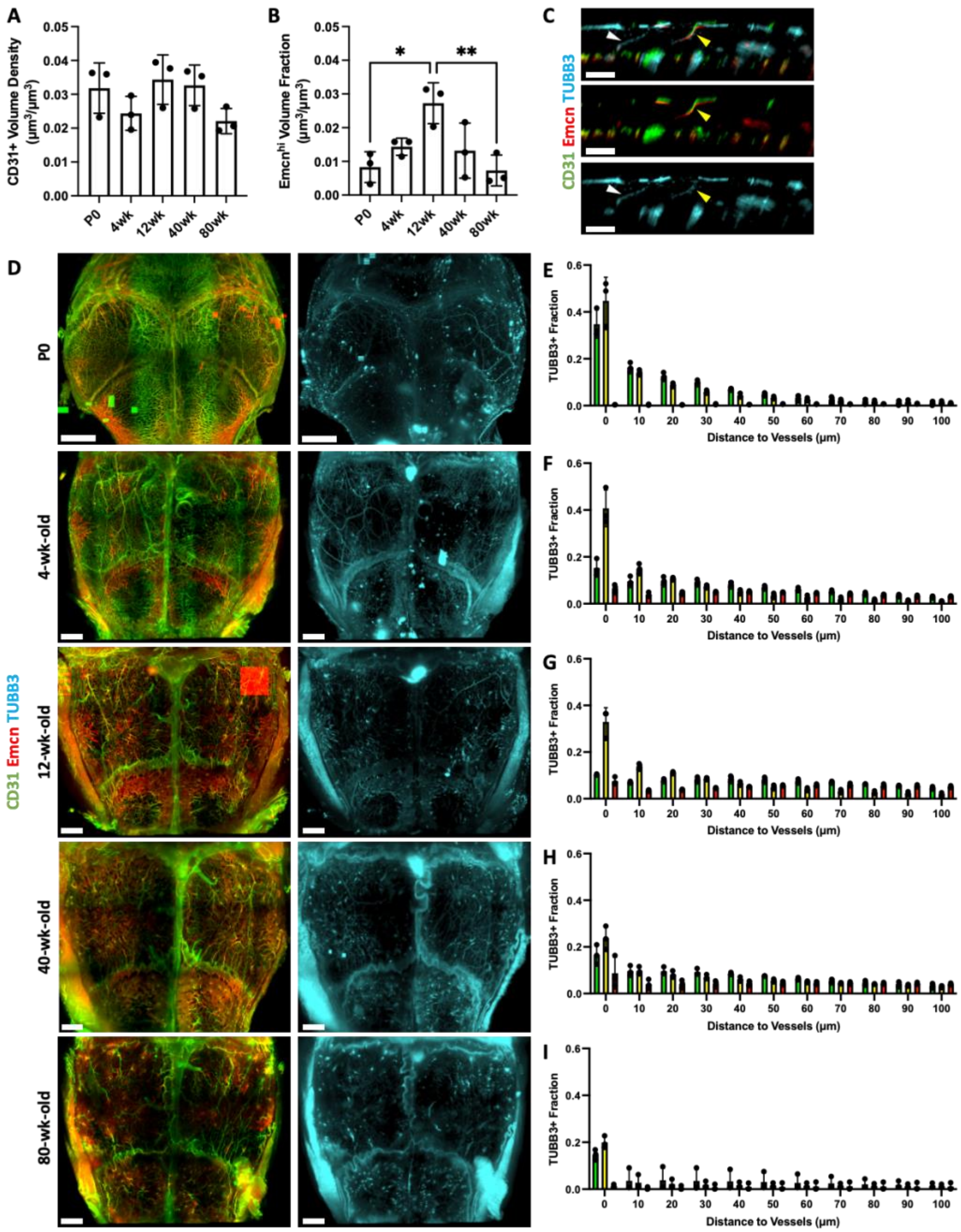


**Supplemental Figure 3: Changes in TUBB3+ nerve interactions with vasculature during aging.** A) Vessel volume calculations for CD31+ blood vessels for P0, 4wk, 12wk, 40wk, and 80wk calvaria. B) Vessel volume calculations for Emcn+ blood vessels for P0, 4wk, 12wk, 40wk, and 80wk calvaria. C) 50 µm coronal cross-section of TUBB3+ nerves and CD31+ and Emcn+ blood vessels. White arrowheads represent nerves and yellow arrowheads indicate vessels in transcortical canals. Scale bar is 300 µm. D) Full calvaria MIP images for CD31+ and Emcn+ blood vessels and TUBB3+ nerves in P0, 4wk, 12wk, 40wk, and 80wk mice. Scale bar is 1000 µm. E-I) Spatial association histograms from TUBB3+ nerve association to $CD31^{hi}Emcn^{-}$ (green), $CD31^{hi}Emcn^{hi}$ (yellow), and $CD31^{lo}Emcn^{hi}$ (red) blood vessels in E) P0, F) 4wk, G) 12wk, H) 40wk, and I) 80wk mice.

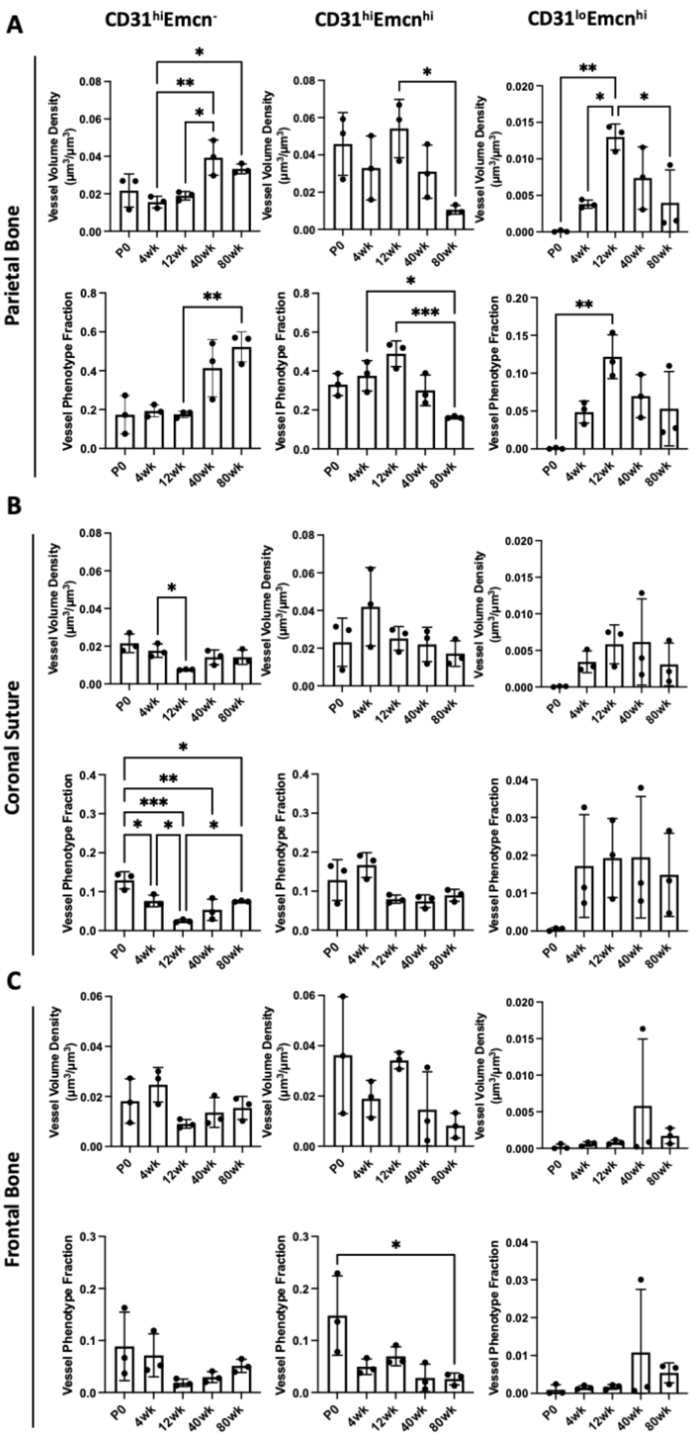


**Supplemental Figure 4: Regional changes in vessel phenotype distributions over the mouse lifespan.** A) Vessel phenotype volume density and fraction for $CD31^{hi}Emcn^{-}$, $CD31^{hi}Emcn^{hi}$, $CD31^{lo}Emcn^{hi}$ blood vessels in the parietal bone for P0, 4wk, 12wk, 40wk, and 80wk calvaria. B) Vessel phenotype volume density and fraction for $CD31^{hi}Emcn^{-}$, $CD31^{hi}Emcn^{hi}$, $CD31^{lo}Emcn^{hi}$ blood vessels in the coronal suture region for P0, 4wk, 12wk, 40wk, and 80wk calvaria. C) Vessel phenotype volume density and fraction for $CD31^{hi}Emcn^{-}$, $CD31^{hi}Emcn^{hi}$, $CD31^{lo}Emcn^{hi}$ blood vessels in the frontal bone for P0, 4wk, 12wk, 40wk, and 80wk calvaria.

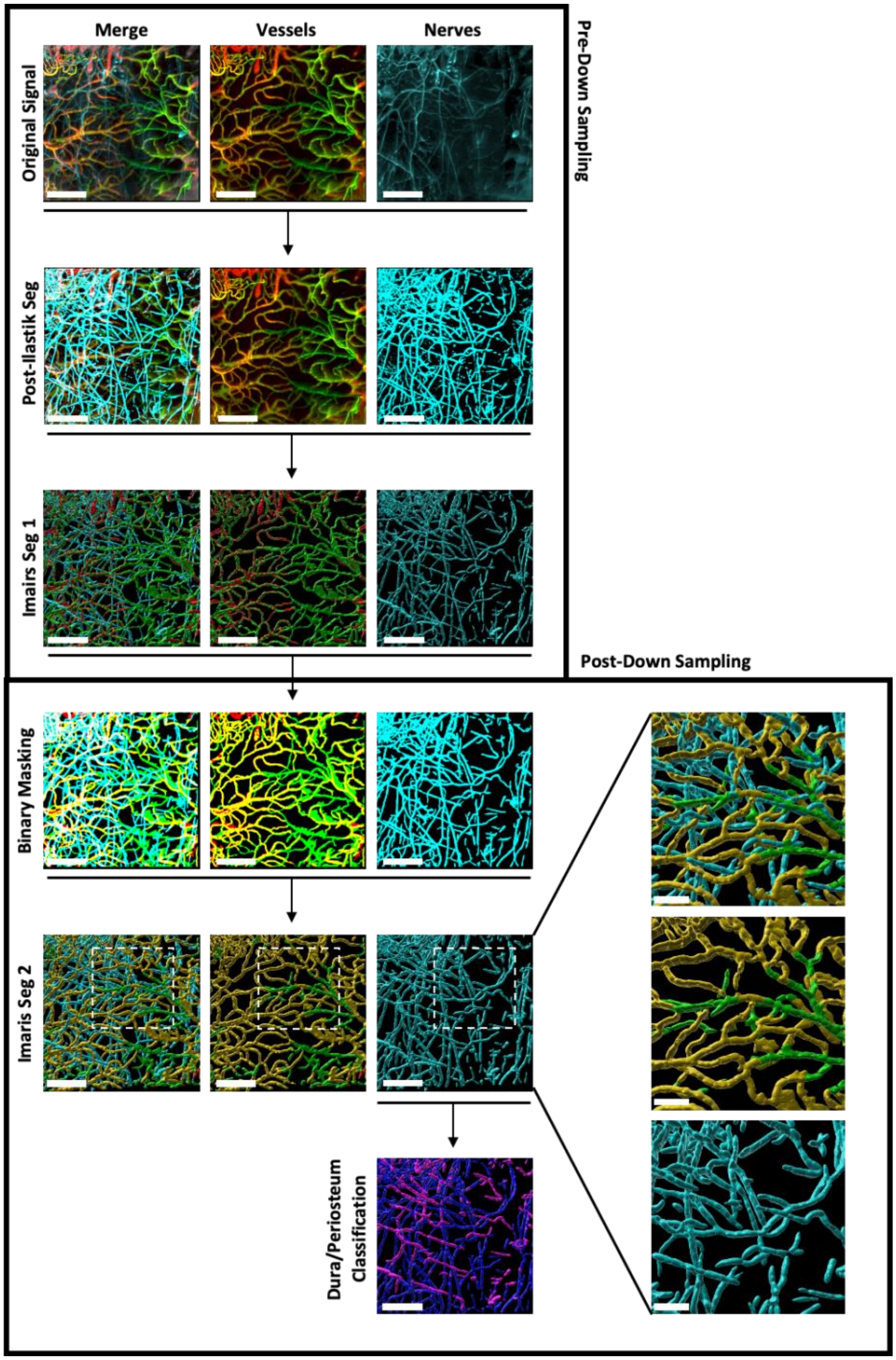


**Supplemental Figure 5: Workflow for image quantification with Imaris®.** Prior to spatial down-sampling and post-Ilastik® nerve segmentation, nerves and blood vessels undergo an initial segmentation with Imaris® to generate individual channel volume data. Following down-sampling, initial Imaris® segmentations are used to generate binary masks for each channel. Next, a secondary segmentation is conducted with Imaris® to generate vessel phenotype classifications, spatial association analysis, and dura/periosteum analysis. The secondary segmentation splits surfaces into 10 µm sections for subsequent analysis (zoomed region). Scale bars are 300 µm. Zoomed region scale bars are 150 µm.

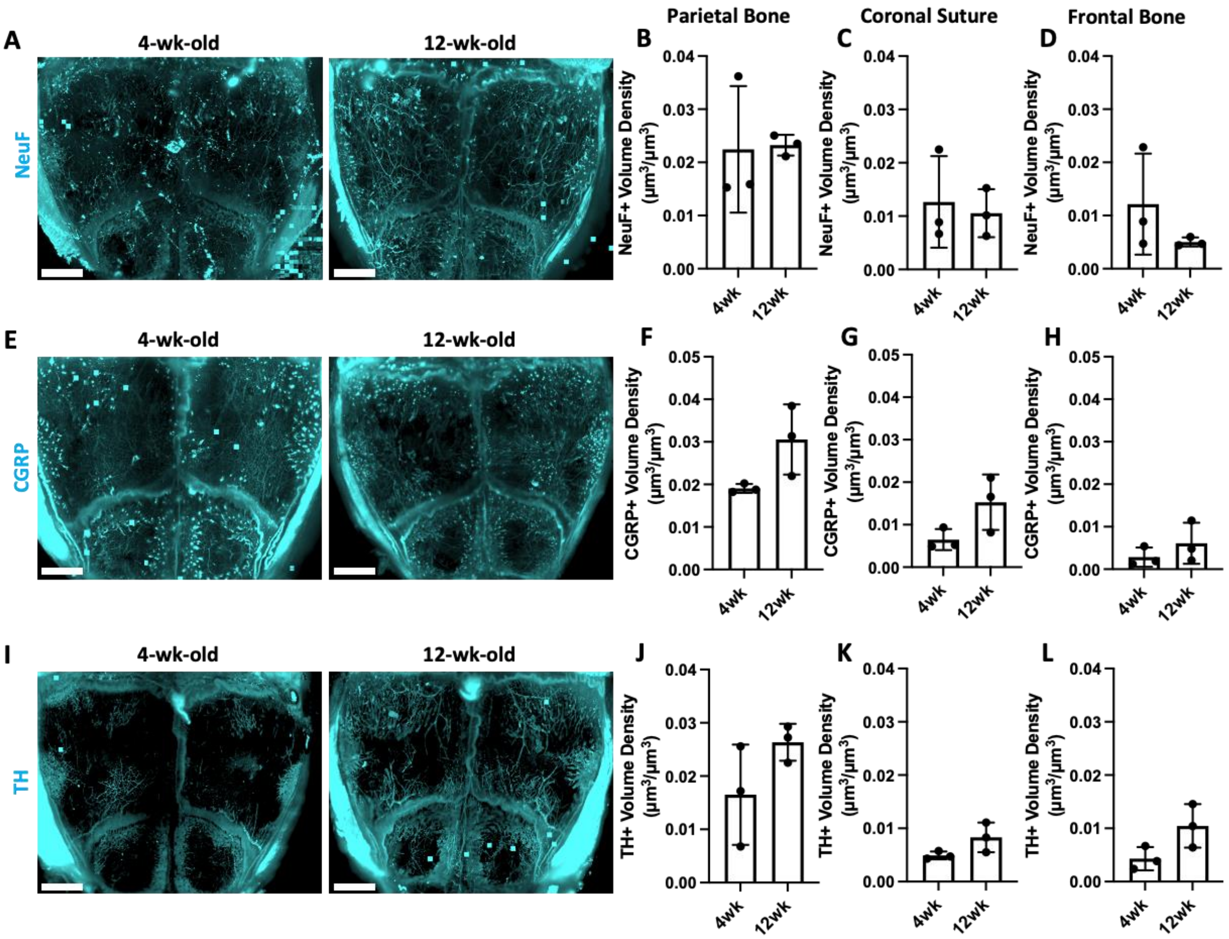


**Supplemental Figure 6: Regional changes in nerve subtype distributions during postnatal development.** A) Full calvaria MIP images of 4wk and 12wk mice with NeuF+ nerves (blue). Scale bar is 1500 µm. B-D) Nerve volume fraction calculations for 4wk and 12wk NeuF+ nerves in the B) Parietal Bone region, C) Coronal Suture region, and D) Frontal Bone region. D) Full calvaria maximum intensity projections of 4wk and 12wk mice with CGRP+ nerves (blue). Scale bar is 300 µm. F-H) Nerve volume fraction calculations for 4wk and 12wk CGRP+ nerves in the F) Parietal Bone region, G) Coronal Suture region, and H) Frontal Bone region. I) Full calvaria maximum intensity projections of 4wk and 12wk mice with TH+ nerves (blue). Scale bar is 300 µm. J-L) Nerve volume fraction calculations for 4wk and 12wk TH+ nerves in the J) Parietal Bone region, K) Coronal Suture region, and L) Frontal Bone region. Data are mean ± SD. Statistics were performed with a two-way ANOVA with post-hoc Tukey HSD test and a two-tailed t-test.

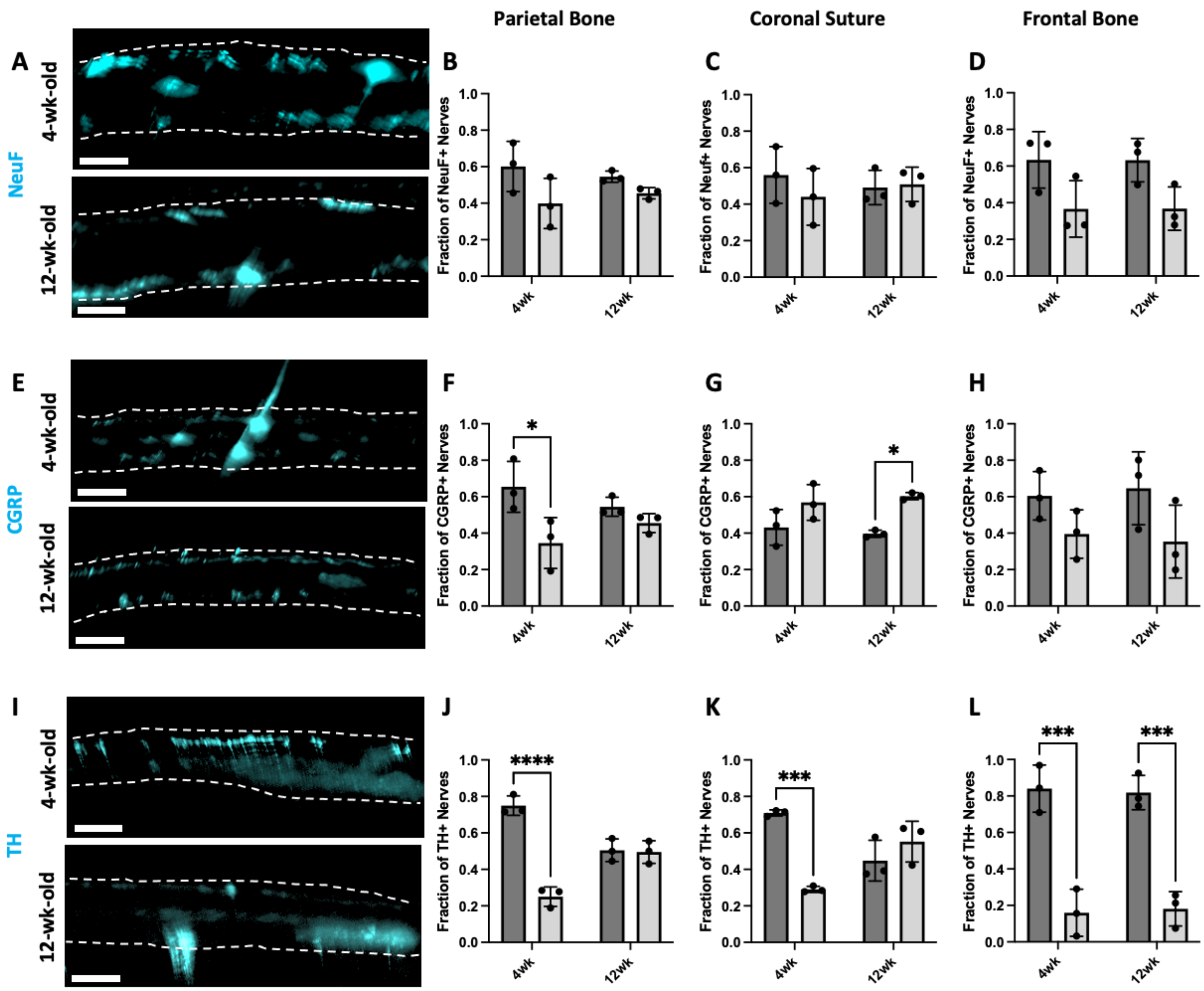


**Supplemental Figure 7: Regional changes in nerve subtype dura and periosteum proportions during postnatal development.** A) 50 µm coronal cross-sections of NeuF+ nerves in 4wk and 12wk mice. Scale bar is 300 µm. B-D) Dura and periosteum calculations for 4wk and 12wk NeuF+ nerves in the B) Parietal Bone region, C) Coronal Suture region, and D) Frontal Bone region. D) 50 µm coronal cross sections of CGRP+ nerves in 4wk and 12wk mice. Scale bar is 300 µm. F-H) Dura and periosteum calculations for 4wk and 12wk CGRP+ nerves in the F) Parietal Bone region, G) Coronal Suture region, and H) Frontal Bone region. I) 50 µm coronal cross sections of TH+ nerves in 4wk and 12wk mice. Scale bar is 300 µm. J-L) Dura and periosteum calculations for 4wk and 12wk TH+ nerves in the J) Parietal Bone region, K) Coronal Suture region, and L) Frontal Bone region. Data are mean ± SD. Statistics were performed with a two-way ANOVA with post-hoc Tukey HSD test and a two-tailed t-test. *$p<0.5$ and ***$p<0.01$ where designated**.**

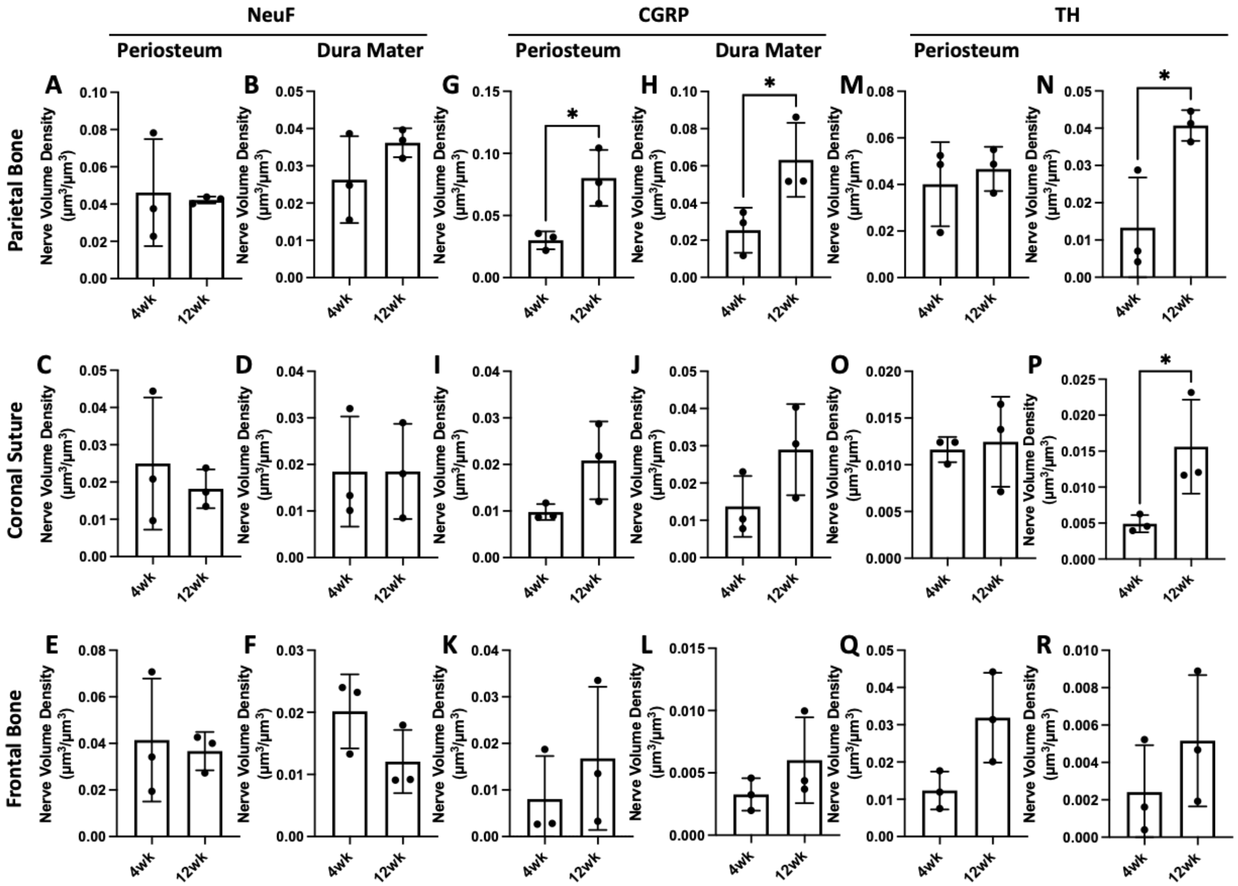


**Supplemental Figure 8: Regional changes in nerve subtype dura and periosteum volume densities during postnatal development.** A-F) Periosteal and dural nerve volume density calculations for NeuF+ nerves in 4wk and 12wk samples in A) Parietal Bone Region Periosteum, B) Parietal Bone Region Dura Mater, C) Coronal Suture Region Periosteum, D) Coronal Suture Region Dura Mater, E) Frontal Bone Region Periosteum, F) Frontal Bone Region Dura Mater. G-L) Periosteal and dural nerve volume density calculations for CGRP+ nerves in 4wk and 12wk samples in G) Parietal Bone Region Periosteum, H) Parietal Bone Region Dura Mater, I) Coronal Suture Region Periosteum, J) Coronal Suture Region Dura Mater, K) Frontal Bone Region Periosteum, L) Frontal Bone Region Dura Mater. M-R) Periosteal and dural nerve volume density calculations for TH+ nerves in 4wk and 12wk samples in M) Parietal Bone Region Periosteum, N) Parietal Bone Region Dura Mater, O) Coronal Suture Region Periosteum, P) Coronal Suture Region Dura Mater, Q) Frontal Bone Region Periosteum, R) Frontal Bone Region Dura Mater. Data are mean ± SD. Statistics were performed with a two-way ANOVA with post-hoc Tukey HSD test. *$p<0.5$ where designated.

**Supplemental Table 1.** List of all key reagents and resources used in this study.

| Reagent or Resource | Source | Identifier |
|---|---|---|
| ***Antibodies*** | | |
| Rabbit anti-mouse TUBB3 (1:200) | Abcam | ab18207 |
| Rabbit anti-mouse NeuF (1:200) | Thermo Fisher Scientific | PA3-16721 |
| Rabbit anti-mouse CGRP (1:200) | Sigma Aldrich | C8198 |
| Rabbit anti-mouse TH (1:200) | Sigma Aldrich | AB152 |
| Goat anti-mouse/rat CD31 (1:200) | R&D Systems | AF3628 |
| Rat anti-mouse/rat Endomucin (1:50) | Santa Cruz Biotechnology | sc-65495 |
| Donkey anti-goat AF800 plus, 0.67 mg/mL (1:100) | Thermo Fisher Scientific | A32930 |
| Donkey anti-rabbit AF647 plus, 0.67 mg/mL (1:300) | Thermo Fisher Scientific | A32795 |
| Donkey anti-rat biotin, 0.75 mg/mL (1:200) | Thermo Fisher Scientific | A18749 |
| Streptavidin AF555 conjugate, 0.67 mg/mL (1:200) | Thermo Fisher Scientific | S32355 |
| ***Reagents*** | | |
| Heparin sodium salt from porcine mucosa | Sigma Aldrich | H3393-50KU |
| Paraformaldehyde, 16% aq. soln., methanol free | Alfa Aesar | 433689M |
| Normal donkey serum | Sigma Aldrich | D9663-10ML |
| Trizma base | Sigma Aldrich | T6066-1KG |
| Trizma hydrochloride | Sigma Aldrich | T5941-1KG |
| Sodium chloride | Sigma Aldrich | S5886 |
| Tween 20 | Sigma Aldrich | P7949 |
| Dimethylsulfoxide | Thermo Fisher Scientific | PI20688 |
